# Mutational signature decomposition with deep neural networks reveals origins of clock-like processes and hypoxia dependencies

**DOI:** 10.1101/2023.12.06.570467

**Authors:** Claudia Serrano Colome, Oleguer Canal Anton, Vladimir Seplyarskiy, Donate Weghorn

## Abstract

DNA mutational processes generate patterns of somatic and germline mutations. A multitude of such mutational processes has been identified and linked to biochemical mechanisms of DNA damage and repair. Cancer genomics relies on these so-called mutational signatures to classify tumours into subtypes, navigate treatment, determine exposure to mutagens, and characterise the origin of individual mutations. Yet, state-of-the-art methods to quantify the contributions of different mutational signatures to a tumour sample frequently fail to detect certain mutational signatures, work well only for a relatively high number of mutations, and do not provide comprehensive error estimates of signature contributions. Here, we present a novel approach to signature decomposition using artificial neural networks that addresses these problems. We show that our approach, SigNet, outperforms existing methods by learning the prior frequencies of signatures and their correlations present in real data. Unlike any other method we tested, SigNet achieves high prediction accuracy even with few mutations. We used this to generate estimates of signature weights for more than 7500 tumours for which only whole-exome sequencing data are available. We then identified systematic differences in signature activity both as a function of epigenetic covariates and over the course of tumour evolution. This allowed us to decipher the origins of signatures SBS3, SBS5 and SBS40. We further discovered novel associations of mutational signatures with hypoxia, including strong positive correlations with the activities of clock-like and defective DNA repair mutational processes. These results provide new insights into the interplay between tumour biology and mutational processes and demonstrate the utility of our novel approach to mutational signature decomposition, a crucial part of cancer genomics studies.

---

Mutations in the genetic material play a fundamental role in the evolution of populations. Their spatial distribution along the genome contains information about the regional variability of the DNA mutation rate and provides insight into the replication timing, gene expression and epigenetic landscape of the underlying tissue [1, 2, 3, 4]. The mutation rate also depends on the DNA sequence context. For single-nucleotide variants, this dependence in a given data sample is usually described using trinucleotide context-dependent relative mutation probabilities, which give the mutational profile of the sample. The observed mutational profile of a sample can be viewed as a linear superposition of different underlying mutational processes. In principle, using a large sample collection, these mutational processes can be identified with mathematical methods such as nonnegative matrix factorisation (NMF), in what is referred to as *de novo* signature extraction. Since this procedure depends on the number of samples available and the variability between samples, the definition of mutational signatures improves with the ever-increasing number of sequenced samples, leading to newer versions of the catalogue of *bona fide* signatures. Many, but by no means all, of the mutational signatures extracted to date have also been associated with biochemical processes of DNA damage or repair. One widely used signature catalogue is provided by the COSMIC database [5, 6].

Once a catalogue of *bona fide* extracted signatures is available, the mutational profile of any sample can be decomposed into the signatures active in the sample without the need for data from an entire cohort. The result of this signature decomposition (also called “signature refitting”) is a list of weights indicating the relative contribution of each signature to the observed mutations in the sample. To our knowledge, all existing methods solve this decomposition problem using a non-negative least squares approach (NNLS) applied to the 96-component mutational profile of the sample. However, we have found that this has several drawbacks:

1. Poor performance for small numbers of mutations. For example, the deconstructSigs method [7] recommends the use of at least 50 mutations, but achieves only limited accuracy even with hundreds of mutations due to the high sampling noise. However, due to the higher cost efficiency, often only whole-exome sequencing (WES) data with mutation counts in the hundreds are available. And even when whole-genome sequencing (WGS) data are available, some cancers have only about 10-100 mutations genome-wide. The application of existing algorithms to subsets of mutations, e.g., from specific parts of the genome or the history of the tumour, is entirely out of scope.
2. Failure to detect certain signatures. Most notably, the single-base substitution (SBS) signature SBS5, a clock-like signature that is present in almost every tumour or normal tissue sample at usually the highest individual signature weight, is often assigned zero weight in all tested methods due to its relatively featureless profile.
3. Lack of error estimates for the signature weights. Even when existing approaches provide error estimates for the derived signature weights, these are determined by bootstrapping the mutations in the sample and not by the accuracy of the weight estimate itself [8].

Here we present an approach to mutational signature decomposition that addresses all of the above problems. Our deep learning algorithm utilises the information contained in the signature frequencies, their covariances, and higher-order dependencies observed in real data. In this way, the method can account for the ubiquity of the clock-like processes underlying SBS1 and SBS5, which occur in varying amounts in almost every sample of both healthy and tumour tissue. On the other hand, it captures strong correlations such as those between the signatures SBS2 and SBS13, both of which are associated with the activity of members of the APOBEC protein family [9]. The increasing resolution of signature extraction methods with increasing amounts of sequencing data also means that previously super-processes can be resolved into finer components, which then occur with very high correlation. An example of this is the initially single UV radiation-associated signature 7, which in the latest versions of the signature catalogue (v3) has been split into the independent but highly correlated components SBS7a, SBS7b, SBS7c and SBS7d. Algorithms that do not take such correlations into account and treat all signatures independently of each other are therefore likely to be at a clear disadvantage. In line with this, we show below that our method, SigNet, outperforms all existing approaches tested, especially for small and medium sample sizes.

The inference of the active mutational processes in a tumour sample provides information about the development of the tumour, the genetic state [10, 11] and the exposure to mutagens [12, 13, 14]. In addition, it enables the classification of tumours into subtypes [15]. Mutational signatures are also markers of therapeutic vulnerability [16, 17, 18] and drug sensitivity [19]. It has also been shown that the activity of mutational processes is heterogeneous across genomic regions, which can be investigated by linking signature-specific mutational activities to known covariates of mutation rate [20, 21]. Finally, a hallmark of tumour development is hypoxia [22], which has been associated with a number of molecular changes in DNA repair [23], which in turn should leave traces in the mutational profile of hypoxic tumours. Recent work reported associations between hypoxia and certain mutational processes in a WGS cohort of 1188 tumours [24], partially consistent with expectations based on experimental data [23]. At the same time, to our knowledge, a much larger WES cohort of over 10000 tumours [25] has not yet been characterised.

Here, we first utilised SigNet’s ability to predict mutational signatures in low mutation count samples to determine signature weights for a WES cohort of over 7500 samples. We then inferred signature activity across tumour history and across functionally distinct genomic regions, revealing pseudo-temporal and epigenetic preferences of DNA damage and repair. Our method allowed us to replace existing approaches to signature attribution in epigenetically marked genomic regions based on genome-wide weight estimates with strictly local prediction. We used this to quantify the dependence of mutational signatures on replication time, histone marks and insertion/deletion (indel) load. With this, we deciphered the mutational processes underlying the ubiquitous but enigmatic signatures SBS5 and SBS40 and locally linked SBS3 to indels and structural variants. Finally, we used SigNet to uncover links between hypoxia and mutational processes that were invisible in a small WGS cohort, reversing some previous findings and revealing a number of new associations with processes involved in DNA replication, DNA repair and basal cellular mutagenesis.

## Results

### SigNet algorithm

Our approach to signature decomposition is based on a deep neural network. The main challenge with such a data-based approach is the need for a labelled data set that can be used as ground truth for training the model. We based our ground truth data set on the SigProfiler assignments of a pan-cancer whole-genome sequencing cohort (PCAWG), available in [26]. We considered this set of samples the best possible set, as it was computed using the algorithm that extracted the COSMIC signatures that form the catalogue of *bona fide* signatures we used. From the combinations of signatures found in the different samples of this set, we created a labelled training data set containing the corresponding mutational inputs with varying total number of mutations. We used these synthetic but realistic data to train the sample-specific mutational profile decomposition tool SigNet Refitter. Given the trinucleotide contexts of a given set of mutations from a sample and the trinucleotide abundances of the sequenced genomic regions, SigNet Refitter outputs the vector of estimated signature weights present in the sample and their error intervals. These are determined in three steps: (1) rough estimation of signature weights from NNLS (only if the number of mutations is large), (2) refined estimation of the signature weights with the Finetuner module, and (3) error estimation of the predicted weights using the ErrorFinder module.

Since the signature composition of a new sample may differ from the training data, we also implemented a sample classifier, called SigNet Detector. This component of the algorithm flags samples that deviate from the expected patterns. These may contain biologically interesting information or signal the presence of mutation calling artefacts (**Extended Data Figure 1** shows the Detector performance as a function of sample size). Since such samples that fall outside the training distribution may exhibit poorer signature decompositions using the neural network, SigNet Refitter instead performs NNLS minimisation without providing error bars for the weight estimates.

Finally, we implemented SigNet Generator, a tool to generate realistic-looking cancer mutation data based on a variational autoencoder (VAE). This tool overcomes the limitation of existing methods of randomly assembling mutational signatures without considering their correlations [27]. Although the more realistic synthetic tumour mutation profiles may be useful for researchers interested in modelling such data, we did not use SigNet Generator to generate training samples for SigNet Refitter because we found that some signature correlations were slightly artificially enhanced by the underlying VAE (**Extended Data Figure 2**). **Figure 1a** shows a diagram of the different modules encapsulated in SigNet. More details on the individual modules can be found in **Methods**.

**Figure 1:**
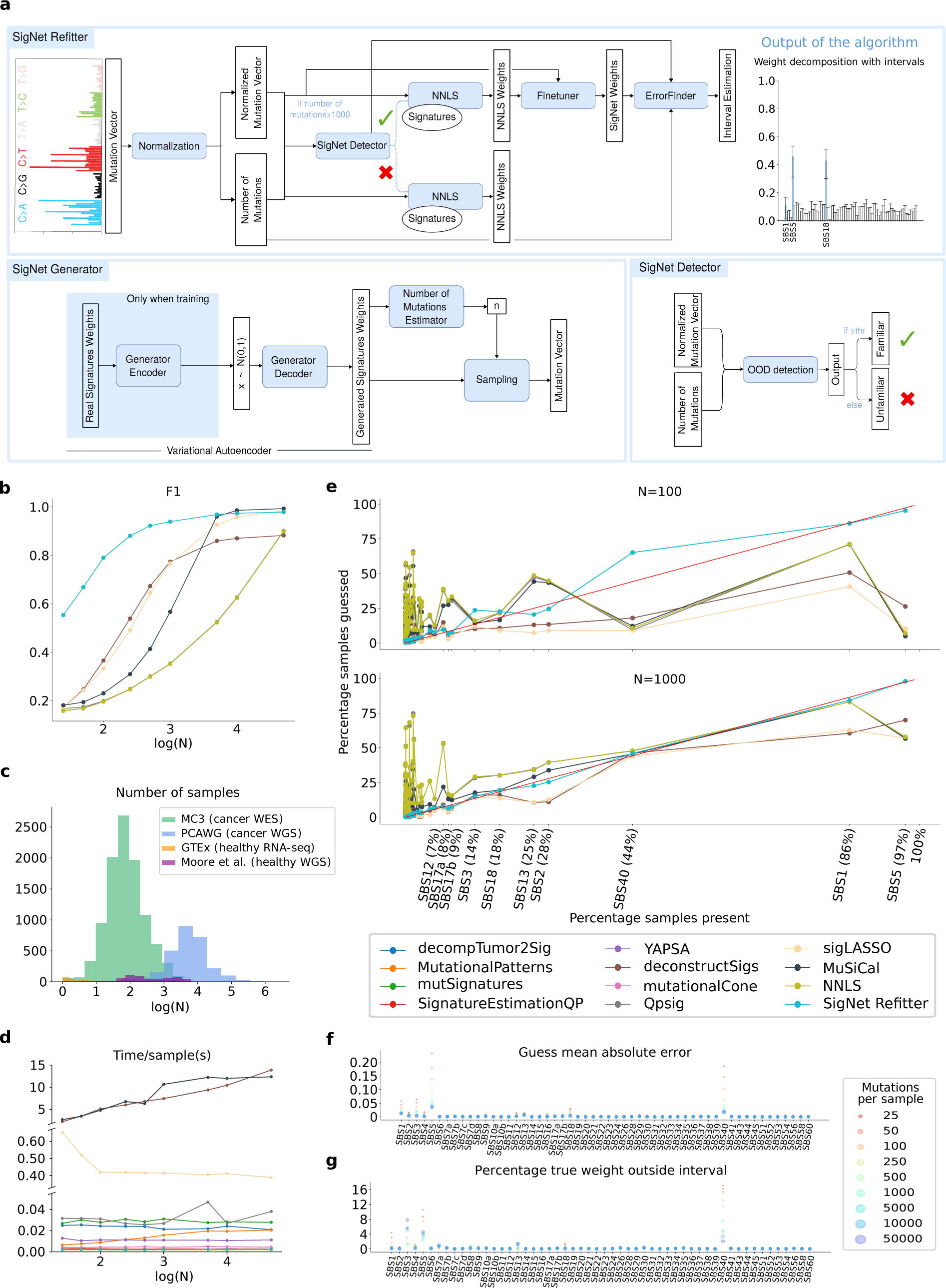
Overview of SigNet and benchmarking against other refitting methods. **a**, Diagram of the different modules in SigNet. Inputs and outputs of all neural networks are shown as white rectangles. **b**, F1 score for SigNet Refitter and eleven other signature refitting algorithms as a function of the number of mutations in the sample, *N* (logarithmic scale). **c**, Distribution of the number of mutations per sample for four different mutation data sets, including from cancer and healthy tissues. **d**, Computation time per sample for each of the different algorithms. **e**, Percentage of samples with a predicted weight of a given signature ≥ 0.01 as a function of the percentage of samples truly containing the signature, for *N* = 100 and *N* = 1000. True weight occurrences taken from PCAWG data. Red line denotes perfect prediction. **f**, Mean absolute error (MAE) between the weight labels and the SigNet Refitter guesses in the test set for each signature and different numbers of mutations. **g**, Uncertainty of the error intervals provided by SigNet Refitter for each signature and different numbers of mutations. This error is computed as the percentage of times that the real value falls outside the error interval. **f** and **g** show signatures that are present with weight larger than zero in at least one sample in the PCAWG data set. WES=whole-exome sequencing, WGS=whole-genome sequencing, NNLS=non-negative least squares.

### Evaluation on synthetic data

We first evaluated the performance of SigNet Refitter on labelled data by using 10-fold cross-validation with the signature assignments obtained from PCAWG [26], preserving the proportions of cancer types and sampling mutation counts from 25 to 10^6^. We compared the performance of SigNet with the state-of-the-art of existing decomposition algorithms, including decompTumour2Sig [28], MutationalPatterns [29], mutSignatures [30], SignatureEstimationQP [31], YAPSA [8], deconstructSigs [7], mutationalCone [32], QPsig [33], sigLASSO [34] and MuSiCal [35]. **Figure 1b** shows the F1 score for varying sample mutation count, covering typical ranges of available tumour mutation data sets (**Figure 1c**). In particular, for samples with less than 1000 mutations, which includes almost all of the more than 10000 WES samples from the MC3 database [25], our method outperforms existing approaches by a wide margin. Interestingly, when comparing all other tested algorithms with each other, we found highly similar performances. This highlights that, despite differences in the details of each implementation, their performance is limited by the NNLS distance minimisation on which all these methods are based. The only methods that show a different performance are deconstructSigs [7], sigLASSO [34] and MuSiCal [35], which is due to the regularisation of signature weights imposed by default. In the case of deconstructSigs, any signature weight smaller than 0.06 is set to zero, while sigLASSO uses L1 regularisation, both resulting in higher accuracy and lower recall as well as longer runtime compared to the rest of the NNLS-based algorithms (**Figure 1d**). MuSiCal also shows better performance compared to the rest of the NNLS algorithms due to the *post-hoc* manipulation of the weights, where signatures are sequentially included and excluded based on the multinomial likelihood of each action. In SigNet, signature weights below 0.01 are set to zero and summarised in the category “Unknown”. While the F1 score (**Figure 1b**) summarises information about precision and recall across all signatures, **Figure 1e** shows the behaviour for different signatures along with their relative occurrence in real data. This shows how difficult it is for NNLS-based methods to accurately predict some of the most common signatures. We also evaluated the mean absolute error (MAE) of the weights output by SigNet Refitter and the accuracy of the error interval as a function of sample mutation count and mutational signatures, **Figure 1f,g**. More properties of the output of the method are given in the **Extended Data**, including the mean error interval length output by ErrorFinder (**Extended Data Figure 3**), the MAE of all the other tested methods (**Extended Data Figure 4**), the full distributions of the error by number of mutations and signature (**Extended Data Figures 5, 6**) and the MAE for all signatures listed in COSMIC v3.1 (**Extended Data Figure 7**).

### Predicting mutational signatures in tumour and healthy tissue

Recognising that much of the currently available sequencing data is WES-based and therefore has a low mutation count, we first derived SigNet predictions for each of 7591 (out of 10218) samples in one of the largest homogeneously curated, publicly available WES dataset, MC3 [25]. Similarly, we decomposed 4728 (out of 4975) samples in the largest homogeneously curated WGS-based metastasis cohort, HMF [36]. The results are summarised in **Figure 2a,b** and all values can be downloaded from **Supplementary Tables 1-4**. To demonstrate the applicability of our method beyond cancer tumour data, we next compared our signature weight predictions with those from a recent study of healthy tissues, including testis [37]. Although being derived from WGS, the number of mutations per sample in this healthy tissue data set is still low compared to WGS data from cancer tumours, ranging from 3 to 6000 mutations (**Figure 1c**). The signature weights in Moore *et al.* [37] were predicted using two *de novo* signature extraction algorithms: the hierarchical Dirichlet process (HDP) and SigProfiler (based on NMF). **Figure 2c** shows the SigNet Refitter weight predictions, correspondingly to Figure 3a from [37], yet with SBS5 and SBS40 shown separately. We found high agreement between the two predictions, suggesting that SigNet Refitter can accurately capture mutational profiles not only of cancer tumours, but also of nonmalignant and germline tissues.

**Figure 2:**
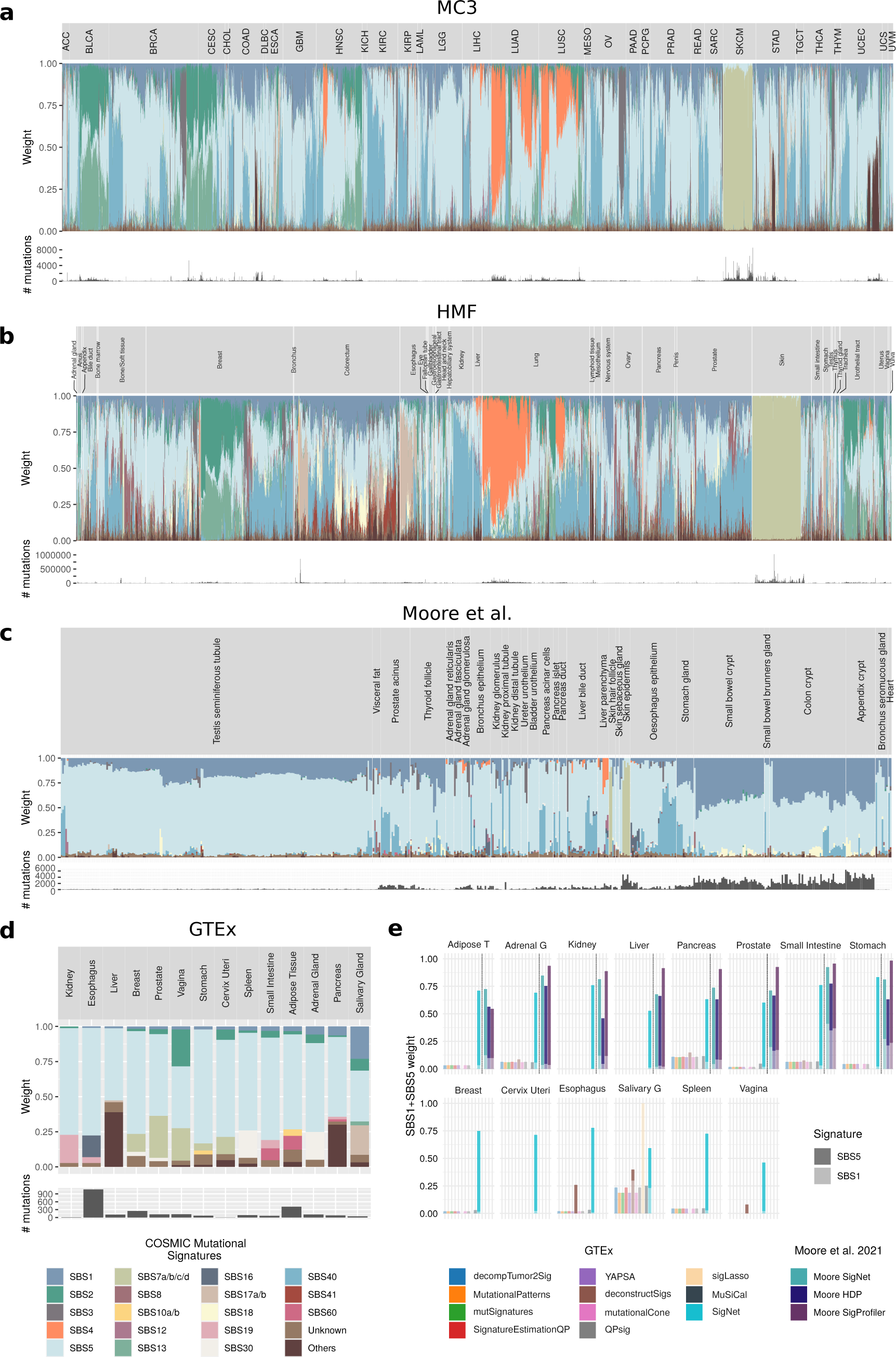
Mutational signature decomposition for real mutation data sets. SigNet Refitter results for **a**, WES data from primary tumours in the MC3 cohort [25], 7591 tumour samples (74%) were classified as “familiar” and are shown. **b**, WGS data from metastatic tumours in the HMF cohort [36], 4728 (95%) “familiar” samples. **c**, WGS data from healthy tissues in Moore *et al.* [37], 536 (96%) “familiar” samples. **d**, RNA-seq data from healthy tissues in GTEx [38]. For each tissue, all available samples were pooled together, 14 (50%) “familiar” tissues. **e**, Comparison of the weight of SBS1 and SBS5 for all the methods benchmarked. First row shows tissues in the GTEx cohort with paired tissue in Moore *et al.* and second row contains GTEx tissues without a pair. Decompositions performed with HDP and SigProfiler in Moore *et al.* are shown for comparison. All the methods except SigNet fail to detect SBS5. “Unknown” corresponds to the sum of all weights lower than 0.01. “Others” combines signatures that have overall low frequency and weight across the different cohorts. Cancer types or tissue names are shown above each plot, number of mutations for each sample below. Within each tissue, samples are displayed according to hierarchical clustering. For cancer type abbreviations, see **Methods**.

**Figure 3:**
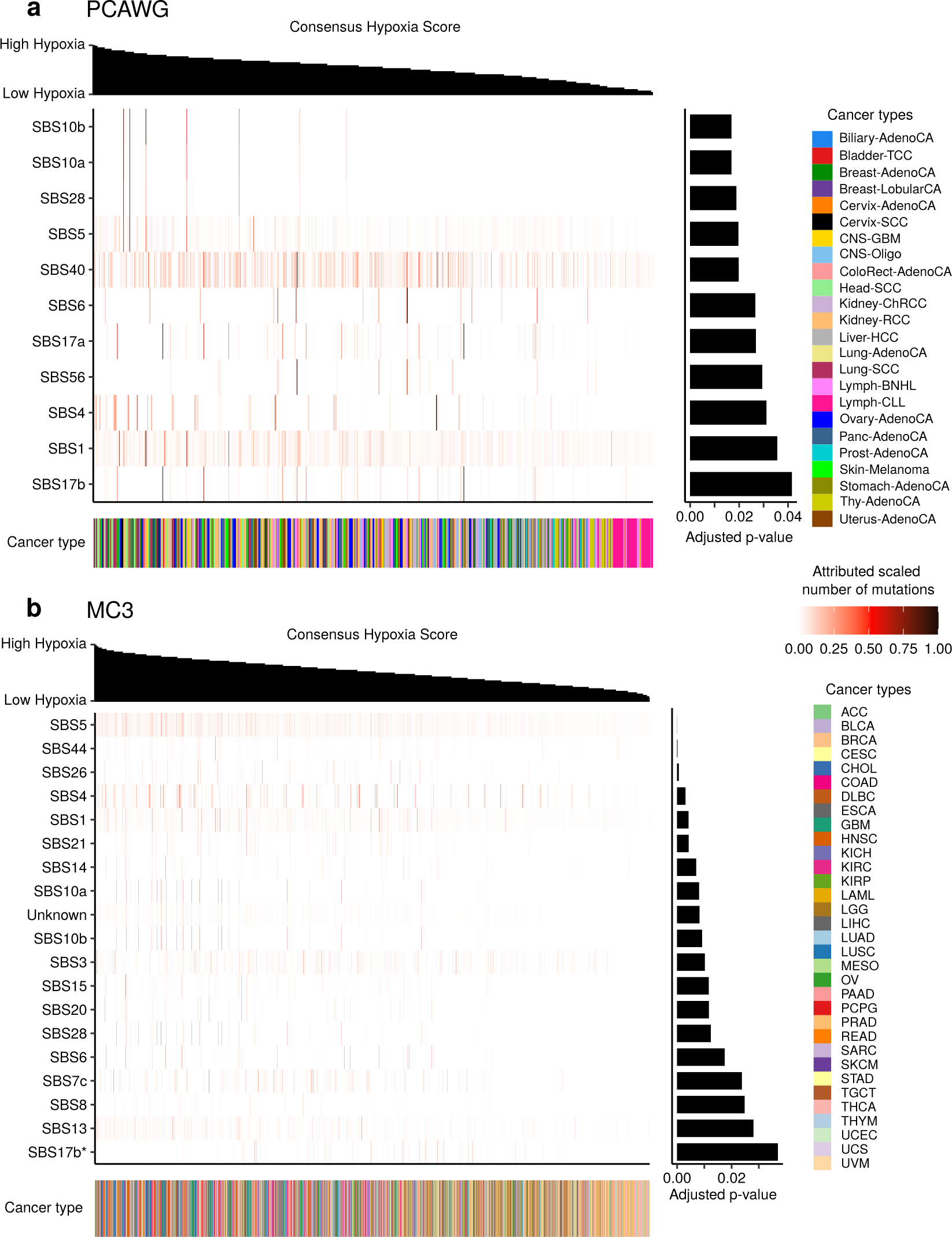
Association between hypoxia and mutational signature activity. **a**, PCAWG data using SigProfiler signature assignments [26] (N=1121). **b**, MC3 data with signature decompositions estimated by SigNet Refitter, restricting to samples that had all explanatory variable information available and were classified as “familiar” (N=7411). Samples are sorted by consensus hypoxia score, signatures are sorted by FDR-adjusted p-value and shown up to *p*_adj_ = 0.05 (black horizontal bars). Significance is based on an ANOVA test, accounting for potential confounding variables, including cancer type (**Methods**). *Negative correlation. For cancer type abbreviations, see **Methods**.

We next applied our method to somatic mutations obtained from RNA sequencing (RNA-seq) of 253 samples of the GTEx cohort [38], pooled in each of the 28 healthy tissues. In principle, mutation calling from RNA-seq is an apt use case for SigNet Refitter, as the problem of low mutation counts is particularly prevalent here (**Figure 1c**). We found that the proportions of the two most common signatures, the endogenous signatures SBS1 and SBS5, in the GTEx cohort were consistent with expectations from experimental data [39], dominating the mutational profile, **Figure 2d**. We then wanted to test how other algorithms would perform on this task. **Figure 2e** shows the comparison of the average weights of SBS1 and SBS5 inferred by the other ten methods and SigNet for the GTEx tissues. We found that all methods except deconstructSigs failed to identify SBS5 in any of the GTEx tissues, while deconstructSigs identified it in 3 of 14 tissues, albeit with low weights. For further validation, we also compared to the healthy-tissue WGS-based predictions for those GTEx tissues that had a match in Moore *et al.* and found them again in very good agreement with the SigNet predictions.

In addition to SBS1 and SBS5, **Figure 2d** also shows several signatures with low biological support identified in the GTEx cohort (e.g., SBS7x in non-sun-exposed tissues). These putatively artefactual signatures occurred even though the abundances of the trinucleotide contexts, i.e., the mutation target, was adjusted to the tissue-specific set of expressed genes (**Methods**) and also in tissues with a mutation count of several hundred. Strikingly, of the six signatures that differed from SBS1, SBS5 and SBS40, five signatures (SBS2, SBS7a, SBS7b, SBS19 and SBS30) had a mutational profile that was almost exclusively restricted to C>T mutations [5]. This is consistent with the excess of such mutations reported in the original study, which was hypothesised to be due to C-to-U RNA modifications [38, 40]. To test this hypothesis, we polarised the mutations attributed to these signatures by transcription strand and found a very strong bias of C>T mutations towards the non-transcribed (coding) strand, which is indeed compatible with an origin of the mutations by RNA-modification (**Extended Data Figure 8**). The last remaining signature identified by SigNet in the GTEx cohort was SBS60, characterised by T>G mutations in the GTG context [5]. Although this type of mutation is relatively common in the DARNED database [41] for exonic RNA modification sites (3% out of all, 10% out of all non-A-to-I modifications), the corresponding mutations neither showed a strong transcription strand bias (**Extended Data Figure 8**), nor did we find an established biochemical aetiology. Therefore, we concluded that they are likely RNA sequencing artefacts, which is consistent with the aetiology reported in COSMIC [5]. In summary, this analysis shows that SigNet can recover the major mutational processes even in very small mutation sets such as those obtained from RNA sequencing, despite a substantial admixture of unknown mutational processes and mutation artefacts.

### Mutational signatures reveal effects of hypoxia on DNA damage and repair

Our signature estimates for the WES cohort (**Figure 2a**) allowed us to investigate the conditions under which different mutational processes occur. One particular condition and a hallmark of cancer is transient or chronic hypoxia, defined as a state of critically low tissue oxygen saturation that is insufficient to maintain adequate homeostasis [42]. Hypoxia has been associated with dramatically increased mutation rates [43, 24], which in turn are attributed to a marked decline in the activity of various DNA repair mechanisms [23]. Since many mutational signatures have been associated with the failure of DNA repair pathways, we hypothesised that a mutational signature decomposition should reveal hypoxia biology if applied to tumours stratified by hypoxia markers.

This idea was also explored in a recent study by Bhandari *et al.*, which found that hypoxia, defined based on gene expression data, decreased SBS1, SBS5 and SBS12, while it increased SBS3, SBS6, SBS17a, SBS17b, SBS21 and SBS56 [24]. To quantify the signatures, the authors used relative signature weights provided by PCAWG [26]. By definition, these weights represent so-called compositional data. This means, for example, that any absolute increase in any one signature must lead to a decrease in the relative weights of all other signatures, even if the absolute contributions of the latter have not changed. Therefore, we hypothesised that this approach may alter the true directions of the correlations under study.

We first repeated the same analysis on the PCAWG data as in Bhandari *et al.* [24], but calculated the correlation between hypoxia and the absolute mutation count associated with each signature (the so-called “activity”) instead of using the signature weights. In contrast to the previous report, this analysis showed that hypoxia actually significantly increased the activities of SBS1 and SBS5 (*p*_adj_ < 0.05), **Figure 3a**. We also discovered a novel positive correlation of hypoxia with SBS40. All three of these processes are clock-like endogenous mutational processes. In the case of SBS1, the source of DNA damage has been linked to the deamination of 5-methylcytosine in a CpG context [9]. This damage is primarily targeted by base excision repair (BER), mediated by the MBD4 and TDG proteins, but it has been proposed that non-canonical DNA mismatch repair (ncMMR) is also involved [44, 45, 46, 47, 48]. Both BER and MMR are suppressed under hypoxic conditions [23], which is supported in our results by an observed correlation of hypoxia with the MMR deficiency-associated SBS6. In the case of SBS5 and SBS40, the causes for both damage and repair are less clear. Recent experimental evidence suggests that one of the possible causes of SBS5 damage may be protonation of single-stranded DNA (ssDNA) [49, 50], while the formation of ssDNA is known to be increased in acute hypoxia [51]. Notably, these experimental studies also found that the source of the damaging protons may be glycolytic sugar metabolisation, which is strongly upregulated under hypoxic conditions [52]. Taken together, these results suggest that the increase we observed in SBS1, SBS5, and thus SBS40 mutation count under hypoxia reflects true tumour biology.

Among the remaining four signatures that showed a novel positive correlation with hypoxia, we surprisingly detected all three signatures associated with mutations of the exonuclease domain of polymerase *ε* (POLE): SBS10a, SBS10b and SBS28. This is particularly intriguing, as it indicates either increased POLE-associated mutagenesis or less efficient repair of POLE-induced mutations under hypoxic conditions. The final novel hypoxia association found in the PCAWG data involves the tobacco smoke-associated SBS4. The link to this signature is likely a direct consequence of the activation of hypoxia-inducible factor 1 by cigarette smoke [53]. In addition, we recovered the positive correlations between hypoxia and SBS17a, SBS17b and SBS56 found in Bhandari *et al.* Of note, to quantify hypoxia the authors selected one from a set of gene expression-based hypoxia scoring schemes, described in [54]. However, we found that the multitude of existing hypoxia scores [55, 54, 56, 57] are not very comparable with each other and therefore derived a consensus hypoxia score (**Methods**, **Supplementary Table 5**). Further, in addition to cancer type, patient age, patient sex and tumour purity, our regression model also controls for average tumour sequencing coverage (**Supplementary Table 6**). **Extended Data Figure 9** shows the results when the scheme of Bhandari *et al.* is retained and only the weights for activities are exchanged.

We then used the ability of SigNet Refitter to fit sparse mutational profiles to estimate the correlations between hypoxia and mutational signatures in the much larger WES data cohort, **Figure 3b** (**Supplementary Table 7**). We recapitulated the positive associations with the activities of SBS1, SBS4, SBS5, SBS10a, SBS10b and SBS28 seen in the PCAWG data set (*p*_adj_ < 0.05). We also found that SBS3, which is closely linked to homologous recombination (HR) repair deficiency (HRD), a process that increases under hypoxic conditions [23], increases with hypoxia [24]. This was confirmed by SBS8, which is associated with SBS3 occurrence in breast cancer tumours with HRD [58]. Among the most remarkable associations we found in the WES cohort were the positive correlations between hypoxia and the activities of signatures SBS6, SBS14, SBS15, SBS20, SBS21, SBS26 and SBS44. These are all seven mutational signatures associated with MMR deficiency [6], which extends the finding of SBS6 in the PCAWG cohort and is consistent with the strong downregulation of MMR genes in hypoxia [23]. Taken together, these results provide strong new evidence for the association of cellular hypoxia with a number of mutational processes that previously had not or inversely been linked to conditions of hypoxia.

### Regional variability and pseudo-temporal shifts of mutational signatures

The activity of mutational processes along the genome is not uniform, but depends on a multitude of epigenetic factors [20, 59]. It is to be expected that this heterogeneity of mutational processes leads to differences in signature activities derived from WES and WGS data. To quantify this, we compared SigNet decompositions of exonic sequences with whole-genome decompositions and with the results from whole genes that contained intronic DNA but excluded intergenic sequences. Both analyses revealed consistent and large genomic context dependencies in mutational signature activities for many signatures (**Figure 4a**).

**Figure 4:**
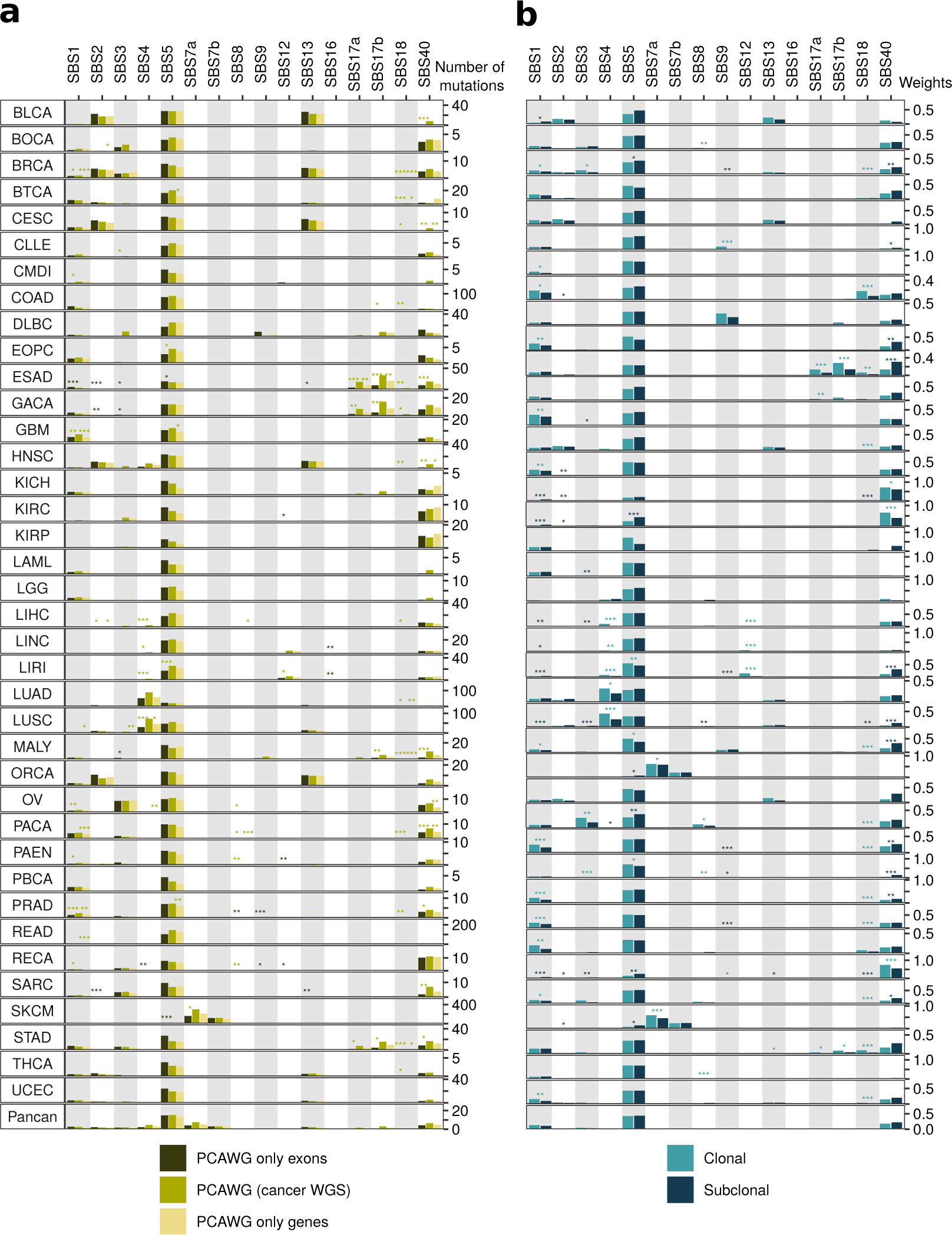
Comparison of mutational signatures between genomic regions and across tumour history. **a**, Mean rescaled number of mutations attributed to each signature across samples for each cancer type for exonic PCAWG mutations (anthracite), all PCAWG mutations (olive) and genic PCAWG mutations (yellow). Only samples classified as “familiar” in the three groups were used (N=1354). The rescaled counts can be converted to whole-genome counts by multiplying by a factor of 286. **b**, Mean mutational signature weights predicted by SigNet Refitter across samples for different cancer types from the PCAWG data set, separated between clonal (turquoise) and subclonal (blue) mutations (N=1767). Note the different y-axis ranges across cancer types. Cancer type names have been mapped to the TCGA study abbreviations. For details, see **Methods**.

To assess the observed differences, we used a recent mapping of mutational signatures to epigenetic states [60], assuming a relative enrichment of the primarily intergenic and intronic part of the whole genome with inactive regions (“quiescent” and “heterochromatin”) compared to exons (“epigenetically active”). For almost all signatures that had a counterpart in Vöhringer *et al.* [60], we found excellent agreement with these assignments: SBS1, SBS4, SBS7a, SBS12, SBS17a/b, SBS18 and SBS40 (corresponding to TS01, TS10, TS05, TS07, TS20, TS18 and TS03, respectively, in [60]) are mainly found in inactive regions and accordingly show a larger mutation contribution in PCAWG compared to that predicted for the exonic data sets. Conversely, SBS2, SBS3, SBS9, SBS13 and SBS16 (TS11, TS19, TS14, TS12 and TS08, respectively) are enriched in active regions, which is consistent with the generally larger predicted mutation contribution in exonic data compared to PCAWG.

In addition to regional variability, another particularly interesting application of tumour mutational profiles is the analysis of pseudo-temporal changes in mutational processes during the course of tumour development. This can be achieved by stratifying mutations according to their cancer cell fraction into clonal (early) and subclonal (late) mutations [61, 62, 63, 64, 65]. However, due to the smaller size of the stratified mutation sets, their signature decompositions are more error-prone. Considering the capabilities of SigNet Refitter in the low mutation count domain, we therefore generated pseudo-time-dependent decompositions of tumour mutational profiles. We note that we could not analyse signature activities here, due to the lack of a proper gauge, and instead report signature weights, which may be subject to compositional effects. Our results, **Figure 4b**, are consistent with previous reports [61, 66], including an enrichment of the APOBEC-associated signature SBS2 among subclonal mutations and clonal occurrence of SBS4 and SBS12. We further found a pronounced clonal trend for the signatures SBS17a, SBS17b and SBS18, which are likely associated with reactive oxygen species (ROS) [67]. Interestingly, this is compatible with the ambiguous role that ROS play over the course of tumour evolution, being beneficial for tumorigenesis, but detrimental for transformed cells [68]. The number of mutations assigned to each signature in **Figure 4a** and the signature weights shown in **Figure 4b** can be found in **Supplementary Tables 8,9**. Taken together, these results suggest that SigNet can help to develop new hypotheses about the biological basis of mutational processes both across genomic regions and over time.

### Possible links between the mutational signatures SBS3, SBS5 and SBS40

In the current COSMIC signature catalogue, three mutational signatures are attributed to clocklike endogenous mutational processes: SBS1, SBS5 and SBS40. However, while SBS1 has been associated with deamination of methylated cytosines in CpG contexts [9], SBS5 and the related SBS40 still lack an aetiology. Various associations with SBS5 have been described, including with mutations in the nucleotide-excision-repair-associated gene *ERCC2* [69]. More recently, evidence was found that SBS5 mutations are independent of replication and instead accumulate due to errors in DNA repair [70]. One possible candidate for the source of DNA damage underlying SBS5 was proposed to be protonation due to glycolysis, specifically targeting ssDNA [50]. Interestingly, we found a strong association of SBS5 and SBS40 with hypoxia (**Figure 3**), a condition in which cells rely on glycolytic sugar metabolism. Support of this hypothesis may also derive from the repeatedly observed correlation between the cigarette smoke-associated SBS4 and SBS5 [13], considering that cigarette smoke is associated with systemic hypoxia that extends beyond the lung tissue [71]. At the same time, about 10000 abasic sites per day are generated by depurination, particularly in ssDNA and partly caused by protonation-induced electron resonance of the N-glycosyl bond [72]. ssDNA arises in many cellular conditions, including replication, transcription, DSBs and uncapped telomeres. *Per se*, ssDNA damage poses a major problem for DNA re-synthesis, as it usually cannot be repaired by dsDNA-dependent repair processes such as BER and NER, thus requiring error-prone translesion synthesis (TLS) or HR-dependent repair.

Both SBS5 and SBS40 have relatively flat mutational profiles, which share features with that of SBS3 (**Figure 5d**), a signature that could therefore shed light on the origins of SBS5 and SBS40. At the sample level, SBS3 was clearly associated with HRD, particularly due to the loss of *BRCA1* or *BRCA2* [73]. This suggests two possible contexts in which the signature might be generated: Replication-associated repair (RAR), due to loss of template switching, and double-strand break (DSB) repair [74]. Recent studies have found synthetic lethality of HRD with loss of polymerase theta (POL*θ*) and upregulation of POL*θ* in HRD tumours [75, 76]. POL*θ*, in turn, is the protagonist of the alternative DSB repair pathway polymerase theta-mediated end joining (TMEJ), which is characterised by excessive generation of ssDNA due to resection [77], but it also plays a role in replication-associated ssDNA repair in HR-deficient cells [78, 79] and can efficiently bypass abasic sites [80]. A key difference between the two repair contexts is the generation of microhomology (MH)-mediated deletions in TMEJ [77], which are not necessarily expected in RAR. Indeed, at the sample level SBS3 has been found to correlate with the indel signature ID6, which is characterized by MH deletions of ≥ 5 bp in length [6].

**Figure 5:**
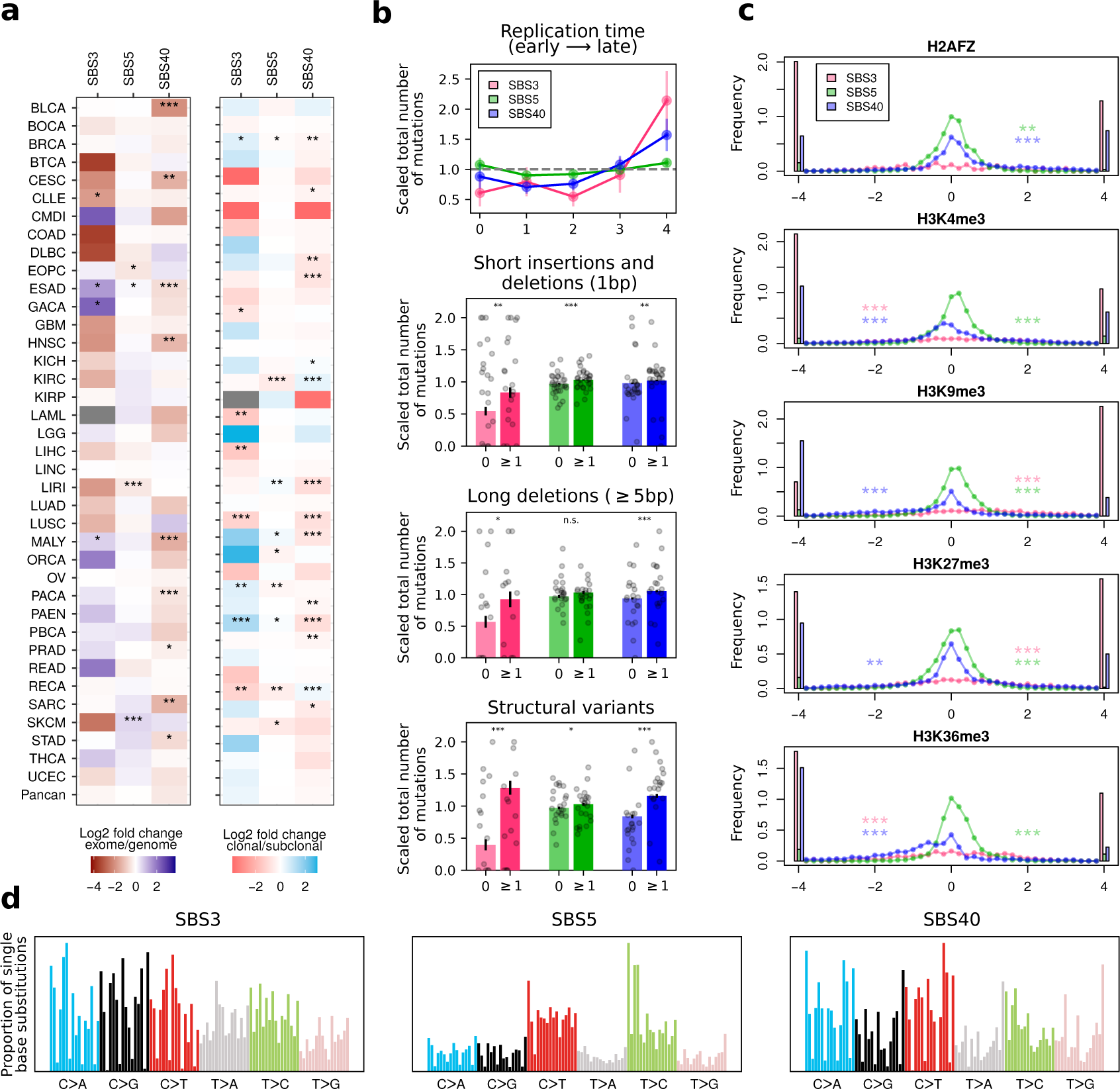
Indel co-localisation and epigenetic dependencies of SBS3, SBS5 and SBS40. **a**, Left: Log2 fold-change of predicted signature activities among PCAWG exonic mutations and all PCAWG mutations. Right: Log2 fold-change of predicted signature weights among PCAWG clonal and subclonal mutations. Pancan = mutations pooled across all tumours. Significance determined with a Brunner-Munzel test. **b**, Top to bottom: Scaled signature activities as a function of replication time (quintiles), presence/absence in nonoverlapping 500-kbp windows of 1-bp indels, deletions of length 5 bp or more, and structural variants. For each signature, the value in each bin was divided by the average value across bins. Top panel: Mean over cancer types. Bottom three panels: Inverse-variance-weighted mean over cancer types that had mutations and were classified as “familiar” in both bins. Error bars denote the standard error of the weighted mean across cancer types. Significance determined with a weighted t-test. **c**, Histograms of log2 fold-changes of signature activities inside and outside of the indicated histone marks, using all experiments in the ENCODE database and correcting for replication time. Lines are linear interpolations between dots. Bars at values -4 and +4 denote instances of zero mutation count inside and outside the histone mark (numerical infinities), respectively. Significance determined with a Wilcoxon rank-sum test and FDR-corrected for multiple testing. **d**, Mutational signature profiles of SBS3, SBS5 and SBS40 [5]. Asterisks denote statistical significance: \**p <* 0.05, \*\**p <* 0.01, \*\*\**p <* 0.001. See **Methods** for details.

To narrow down the source of SBS3 to RAR or TMEJ, we first determined the degree of colocalisation of SBS3 with indels, including long deletions ≥ 5 bp, and structural variants (SVs; **Methods**). We found significant positive associations between SBS3 mutation activity and short indels, long deletions as well as SVs, **Figure 5b**. We then analysed the enrichment of SBS3 in histone marks, correcting for replication time as a putative confounder (**Methods**, **Supplementary Table 10**). Note that SigNet allows direct prediction of signatures in regions with and without the mark, which is expected to be more accurate compared to more indirect tests based on mutation assignment using genome-wide signature weight estimates. This is because genome-wide signature weight estimates are averages that lead to an under-assignment of mutations in regions with high activity and an over-assignment in regions with low activity when there is heterogeneity in the spatial distribution of signature activity. After Bonferroni correction, we found that SBS3 is significantly enriched in H3K9me3 and H3K27me3 (*p*_adj_ <0.01), which are respectively associated with DSB repair via HR (HRR) and TMEJ [81], **Figure 5c**. Conversely, SBS3 is depleted from H3K4me3, H3K27ac, H3K4me1, H3K36me3, H3K9ac, H3K79me2 and H4K20me1, histone marks associated with active genes [82] (**Supplementary Table 11**). These results suggest that SBS3 arises primarily in the context of substitution of HRR by TMEJ, rather than RAR, consistent with the absence of replication strand bias in SBS3 [6]. At the same time, a contribution from ssDNA gap filling during RAR mediated by TLS is expected [78].

Given the likely association between POL*θ* and SBS3, we next asked if SBS40, which is very similar to SBS3, might also be affected by a similar repair process. SBS40 also shows a significant enrichment in regions with short indels, consistent with its reported association with the indel signature ID5 [6]. Furthermore, like SBS3, SBS40 tends to be co-localised with long deletions and SVs, albeit with smaller effect sizes (**Figure 5b**). While we found that SBS40 was significantly depleted at largely the same active gene histone marks as SBS3, the signature was also depleted at the HRR-associated H3K9me3 mark (*p*_adj_ <0.01), suggesting that SBS40 does not occur where functional HRR is active. Only one of the 31 histone marks tested is significantly enriched in SBS40: H2AFZ. Intriguingly, this histone variant is transiently exchanged onto chromatin at DSBs, facilitating both HRR and classical non-homologous end joining (cNHEJ), and failure to remove H2AFZ leads to alternative NHEJ (TMEJ) [83, 84]. This would place SBS40 in the context of DSB repair via TMEJ also in cells with functional cNHEJ and HRR. One possible explanation for this putative association could be specific DSB events in which loading of Ku, the initial DSB end binding factor in cNHEJ, is foreseen but impaired, including end structures with extended ssDNA tails expected after aborted HR or collapsed replication forks. TMEJ has been suggested to be critical in these circumstances, consistent with its role as a crucial alternative to cNHEJ in mammalian cells [77]. Moreover, POL*θ*-mediated repair was reported to be the only pathway capable of repairing replication-associated DSBs resulting from inheritable ssDNA gaps refractory to HRR [85, 86]. The frequency of such events is expected to correlate with increased exposure to DNA damage, which in turn is compatible with the notable tendency of SBS40 to produce late-occurring mutations, as well as its rarity in the human germline (**Figures 5a, 2c**). Finally, the significant enrichment of SBS40 on intergenic regions in a large number of cancer types (**Figures 4a, 5a**) may reflect the requirement for TMEJ at uncapped telomeres and DSBs within telomeric repeats [87].

The final signature that shares similarities with SBS3 and SBS40 is SBS5. Repair of SBS5-causing damage has been proposed to be related to POL*ζ*-mediated TLS in a yeast model and reduced SBS5 activities were found in human breast cancer cell lines lacking the TLS mediator REV1 [49, 88]. Of note, yeast does not possess a POL*θ* orthologue and POL*θ* performs TLS [89]. To clarify whether POL*θ* might play a role in the generation of SBS5, we determined the correlation between sample-specific POL*θ* expression and predicted SBS5 activity in cancer tumours. We found a highly significant correlation coefficient (*p <* 10*^−^*^59^), which was further significantly higher compared to the distribution of correlation coefficients with SBS5 activity across all genes’ expression levels (*p* = 0.005; **Extended Data Figure 10**). A similar trend was observed for SBS3 and SBS40, but with lower statistical significance (*p ≃* 0.1). The correlation between SBS5 and POL*θ* expression may point to competitive binding of POL*θ* and RAD51 during RAR and HRR as the driver of SBS5 [75, 76], especially in early-replicating regions (**Figure 5b**). Finally, we found that SBS5 is enriched in 15 histone marks associated with active promoters, enhancers or gene bodies (*p*_adj_ < 0.01), including eight acetylation-driven marks (**Supplementary Table 11**). Thus, overall, SBS5 may result in part from RAR and DSB repair under benign cellular conditions. The SigNet decompositions for the five different replication time bins can be found in **Supplementary Table 12**, and the weights attributed to the three focal signatures in the windows with short indels, long deletions and SVs can be found in **Supplementary Tables 13-15**. The results for all signatures and all cancer types are shown in **Extended Data Figures 11-16**.

We also wanted to test whether SBS5 or any other signature activity was associated with mutations in specific genes. Any measurement of such an association needs to take into account the intrinsic correlation between the signature activity and the probability of seeing a gene mutated. SBS5 had previously been associated with mutations in *ERCC2*, based on an algorithm developed by Strona *et al.* for ecological applications [90, 69]. An implicit assumption of this algorithm is the equiprobability of mutations across genes or samples, which, however, is not necessarily fulfilled in the context of cancer genomics. Using simulations and a minimal example, we found that this can lead to spurious associations of signature activity with highly mutated genes, including long genes, genes with a high background mutation rate, and cancer driver genes (**Extended Data Figure 17**, **Methods**). Consistent with this, when applying the same algorithm to our data, we found an enrichment of associations of signatures with known cancer drivers and long genes (e.g., *TTN*), which we therefore do not report. Taken together, our results shed light on the genomic distribution and aetiology of three of the most frequent mutational signatures occurring in cancer tumours and in the human germline.

## Discussion

The decomposition of mutational signatures has become an essential part of tumour analysis. Unless new mutational processes are suspected in a given data set and the data set is large, reliable signature decomposition is key. It circumvents the problems of *de novo* mutational signature extraction, such as the need for many and varied samples and the intricate mapping of extracted signatures to *bona fide* signature catalogues, which is often based on suboptimal similarity measures like cosine similarity. With SigNet, we have introduced a new signature decomposition algorithm that enables researchers studying cancer genomics and mutational mechanisms in other biological systems to quickly and accurately estimate the activities of mutational processes. We have demonstrated that machine learning significantly improves our capacity to infer signature contributions in samples with low mutation counts. This opens up the possibility to reliably study samples from cohorts with whole-exome sequencing, cohorts with whole-genome sequencing of healthy tissue, cancers with low mutation rates and subsets of mutations from specific genomic regions, with potential future applications also to more mutation sets from the even sparser RNA or targeted sequencing. Importantly, our tool, SigNet, is easy and fast to run (**Figure 1d**), can be customised to any genomic target, and provides information beyond signature weights: It measures the similarity of the provided sample to the training data (SigNet Detector) and estimates the reliability of the obtained weights by outputting error bars (ErrorFinder module of SigNet Refitter). Once the next version of the catalogue of *bona fide* mutational signatures is released, SigNet can be easily retrained and updated.

Even though trained on WGS data, we have shown that SigNet readily extends to WES data. This allowed us to provide signature decompositions for the largest publicly available and homogeneously curated WES data set [25]. We identified strong novel links between tumour hypoxia and several DNA repair-associated mutational signatures, including all known MMR-deficiency-related (SBS6, SBS14, SBS15, SBS20, SBS21, SBS26 and SBS44), all POLE-related (SBS10a, SBS10b and SBS28) and all known HRD-related single base substitution signatures (SBS3 and SBS8). In addition, we were able to show that SBS1, SBS5 and SBS40, three endogenous and quasi-ubiquitous signatures, also correlate positively with hypoxia, in contrast to previous reports.

SigNet allowed us to compare signature activities across epigenetic features (genomic regions, histone marks, replication time, indel density) in a strictly local manner, i.e., by decomposing the mutational profile of mutations collected in specific regions. This revealed novel associations between epigenetic states and mutational signature activities and allowed us to investigate the origins of SBS3, SBS5 and SBS40, focusing on the links to POL*θ*. The generation of point mutations by POL*θ* has been proposed to be related to its translesion synthesis activity during ssDNA gap filling [89, 78, 79], although other translesion polymerases may also play a role [91, 49, 92, 79]. The increased importance of POL*θ* in both replication-associated and DSB repair in cancer tumours coincides with the increase in replication stress caused by oncogenes [93]. Protons have been proposed as a source of damage underlying SBS5 [50]. While this was shown using an experimental setup involving ssDNA, it is interesting to note that protonation damage has little effect on methylated cytosine in double-stranded form [94]. Coincidentally, depletion of the CpG mutation category is a prominent feature of all three signatures SBS3, SBS5 and SBS40. Further, HR-deficient cells and cells with down-regulated RAD51 were found to be markedly hypersensitive to proton irradiation compared to wild-type cells, and more so than to photon irradiation, whereas the absence of cNHEJ does not result in hypersensitivity to proton irradiation [95]. Considering the role of POL*θ* in the synthetic lethality of PARP1 inhibitors in HRD tumours [96], our results may serve to inform future concepts for cancer therapy.

## Methods

### SigNet Detector

The SigNet Detector artificial neural network (ANN) [97] is in charge of distinguishing realistic-looking samples. Therefore, we perform a binary classification, flagging as 1 those samples that look familiar, and 0 those that do not. To train it, we used binary cross-entropy loss:

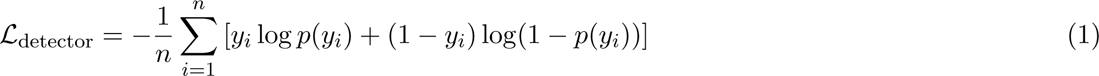

where *y_i_* is the label of sample *i ∈ {*1*, …, n}* (1 for familiar and 0 for unfamiliar) and *p*(*y_i_*) is the predicted probability of the sample being familiar.

The network’s architecture is a feed-forward ANN composed of standard linear layers with leaky ReLU activation [98], and the optimiser used is Adam [99]. The input of the neural network is the 96-element mutation vector containing the total number of mutations for each mutation category, which we decompose into two objects: the total number of mutations (the sum of all the elements in the mutation vector), and the normalised mutation vector (the same vector but dividing it by the total number of mutations). The input vector is normalised so that all the inputs from different samples have the same order of magnitude and the neural network can be trained with more ease. At the same time, it is important to keep the information on the number of mutations, since for low numbers of mutations the samples will be very noisy and the model needs to learn how to overcome this extra layer of complexity. The output of the SigNet Detector is a value between 0 and 1 that represents the probability of the input sample belonging to the familiar class.

The training set is composed of the same number of realistic-looking samples and random samples. The realistic-looking samples were generated by sampling from the real linear combination of signatures provided by SigProfiler in the PCAWG data set [26]. The random set was generated by (1) selecting a random number of signatures (between 1 and 10) and (2) assigning uniform random weights to each, such that each signature has the same probability to be chosen. We generated samples ranging from 25 mutations to the order of 10^4^ mutations. For values larger than that, we used the real distribution without sampling. This ensures that the algorithm is robust to any real sample that the user can provide.

### SigNet Refitter

#### Non-negative least-squares

In mathematical optimisation, the problem of non-negative least squares (NNLS) is a type of constrained least-squares problem, in which the coefficients are not allowed to become negative [100]. In the studied case, this ensures that the solutions are biologically feasible. The mathematical formulation of this problem is the following:

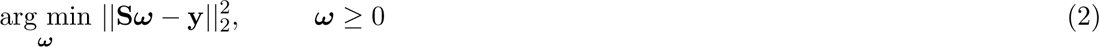

where **S** is a matrix containing the signatures in columns, **y** is a given sample profile (96 mutation counts), and ***ω*** is the vector of weights that we want to find. Here ***ω*** ≥ 0 means that each component or weight must be non-negative, and *|| · ||*_2_ denotes the Euclidean norm.

Therefore, by applying the NNLS optimisation we are minimising the Euclidean distance between a given sample profile and the reconstructed profile. This is what other algorithms use to find the weights of the refitting. However, when the number of mutations in the sample is low, such as in WES, healthy somatic tissue sequencing, RNA sequencing data or even WGS for some cancer types, the input profile does not look like the real profile that corresponds to the linear combination of signatures, due to the high degree of stochasticity. Thus, we hypothesised that this kind of classical algorithm cannot be used in the regime of few mutations and we turned to using artificial neural networks instead.

#### Finetuner

The Finetuner is the neural network of SigNet Refitter in charge of finding the proper signature refitting. For this, we do not have a single neural network but, instead, we have two neural networks for two different regimes: “low” and “high” number of mutations. We make this distinction since low-mutation-count profiles are dominated by noise and the reconstructed profiles might not look like the input profiles. Here, correlations between signatures are the most important feature that the neural network can use to predict the signature weights. However, for high numbers of mutations the reconstructed profiles should be close to the inputs (i.e. NNLS should provide an acceptable initial guess) and this can be used by the neural network to refit the samples.

The input of each neural network is different. For the large number of mutations network we input three different objects: we provide the normalised mutation vector, the number of mutations of the sample, and the baseline guess that we obtained from the NNLS step. The latter element functions as a starting guess for the neural network that should be fine-tuned to find a better fitting. The low number of mutations neural network gets as input the normalised mutation vector and the number of mutations. The NNLS guess is not helpful in this regime. The output of both neural networks is the same: the signature fitting of the sample, which corresponds to a vector of length 72 that contains the weight of each of the signatures in COSMIC v3.1. Note that the signatures that are not present in any of the PCAWG training samples will most likely be learnt as never present. Therefore, the Finetuner part of the algorithm will tend not to include them.

The training set for this neural network is composed of realistic-looking samples generated by sampling from the real linear combination of signatures extracted from the PCAWG data set provided in [26]. For each PCAWG sample, we generated ten samples with different numbers of mutations ranging from 25 mutations to the order of 10^4^.

The Finetuner loss that we used to train the neural network is based on the distance of two categorical probability distributions of signatures, the Kullback-Leibler divergence:

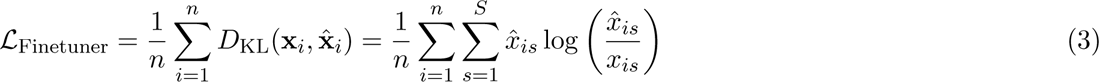

Where 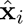 is the predicted weight decomposition across *S* signatures for sample *i*, with *x_is_ ∈* [0, 1] and 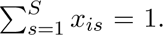 The sample label, **x***_i_*, corresponds to the real weights, and *n* is the batch size. This distance measure is typically used in neural networks of this kind.

#### ErrorFinder

The weight estimates that we find after the Finetuner step are not exact. Their accuracy will depend on both the number of mutations of the sample and the signatures that are present. Namely, when the number of mutations is low, the sample mutational profile has higher sampling noise and it is more difficult to fit than that of a sample with high mutation count. Therefore, we would expect to have a less accurate weight refitting for samples with lower than with higher mutation count. Furthermore, there are some signatures which, because of their shape (i.e. they are flat and do not have specific traits that make them distinguishable), can be more difficult to identify, implying that their weight predictions are less accurate. We want to be able to quantify these different sources of error by finding prediction intervals with a certain confidence level per signature and sample. As far as we are aware, there is little known about prediction intervals for neural networks. We first tried to apply the same architecture of the only article we found that discusses this topic [101]. However, training our network based on that approach failed and we instead developed a modified method that would work for our inference.

In order to find prediction intervals there are two quantities that need to be minimised at the same time: the width of the interval and the distance between the real value and the interval boundaries. The former is necessary so that the intervals have a reasonable width. If we do not impose this, the neural network learns to create intervals that cover the whole range between 0 and 1, making the intervals uninformative. The latter is necessary in order to ensure that some proportion of the real values fall inside the interval. We want them to be as small as possible while containing most of the real values inside of their range (ideally at least 95% in order to have this level of confidence). The corresponding loss function is:

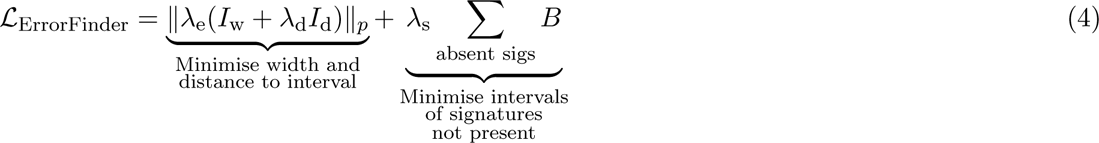

where *I*_w_ is the interval width, *I*_d_ the distance to the interval boundary (set to zero if the real value is inside the interval boundaries), *B* is the upper bound of the interval and *λ*_e_, *λ*_d_ and *λ*_s_ are Lagrange multipliers that allow us to control for the effect of each term in the function. ∥ · ∥*_p_* denotes the p-norm, for which we choose *p* = 5. By tuning the Lagrange multiplier *λ*_d_, we can prioritise the minimisation of *I*_d_ (for large numbers of *λ*_d_) or the minimisation of *I*_w_. In the first case, the precision of the intervals is increased but the width will also be larger. In the second case, the width will decrease but the accuracy will be lower. Since there are signatures that have larger error than the rest because of their flat shape (mainly SBS3, SBS5, SBS40), we decided to set a larger Lagrange multiplier for them in order to force them to have lower error by increasing the width of their intervals. Choosing *λ*_d_ = 0.1 for all the signatures except the ones with larger error where we set it to 0.5, we find a nice balance where the intervals are small enough while ensuring a high level of confidence (see **Figure 1f**). The rest of the parameters were chosen using a Bayesian optimisation of the parameters and were set to *λ*_e_ = 33434 and *λ*_s_ = 0.276.

### SigNet Generator

The SigNet Generator is composed of three modules sequentially called. First, a variational autoencoder (VAE) [102] capable of generating realistic-looking signature compositions. Next, a neural network that estimates the number of mutations that the sample should have, based on the signatures involved. And finally, given the signature composition vector and the number of mutations obtained from the previous two steps, there is a sampling step that generates the corresponding mutational vector.

Variational auto-encoders are deep generative models composed of two blocks: an encoder and a decoder. The encoder non-linearly projects the inputs into a latent space of a pre-fixed dimension, while the decoder recovers the original input from this projection. When sequentially applying encoder and decoder, the function one obtains is close to the identity function. What makes them particularly useful in our case is the application of some conditions in the latent space. For instance, we force the projected data to follow a p-dimensional Gaussian *N* (0, 1) distribution. Therefore, we obtain a mapping between the original data distribution and a standard multidimensional Gaussian. This allows us to generate realistic-looking mutational vectors, simply by sampling from a Gaussian and serving those samples as inputs to the decoder. Effectively, this network is learning the underlying correlations between signatures, being capable of generating previously unseen but plausible examples.

In our case, both the encoder and the decoder are modelled by feed-forward ANNs, composed of standard linear layers with leaky ReLU activation. The module was trained with the label PCAWG data set [26]. Furthermore, we use the re-parametrisation trick [103] for variance reduction: while training, given an input *x_i_*from the data set *D* = *{x_i_}_i_*_=1:*n*_, where *n* denotes the training data set size, the encoder returns two values: *µ_i_*and *σ_i_*. These values are then used to sample the input from the decoder *z_i_ ∼ N* (*µ_i_, σ_i_*), where *N* (*µ, σ*) denotes the normal distribution with mean *µ* and standard deviation *σ*. Finally, the decoder takes this sample *z_i_* and returns a reconstruction *x*^*_i_*. We used the Adam optimiser and the loss function is:

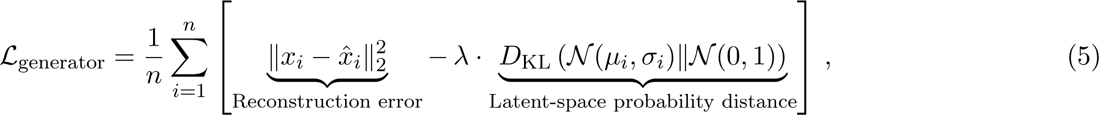

Where *λ* is a Lagrange multiplier balancing both terms. In this case it decreases as training evolves for stability purposes. Further details of the implementation will be available in our GitHub repository (https://github.com/weghornlab/SigNet).

Feeding values from a standardised normal, i.e. *z_i_ ∼ N* (0, 1), to the decoder returns realistic-looking signature combinations *x*^*_i_*. This is one of the outputs of SigNet Generator. The second output is a sample drawn from the obtained combination of signatures. For this, the user can choose whether a realistic number of mutations should be estimated, and if this is the case, said number will be estimated using the second block of SigNet Generator: the number of mutations estimator.

The number of mutations estimator is another feed-forward artificial neural network composed of standard linear layers with leaky ReLU activation [98] and Adam optimiser [99]. The input of the neural network is the vector of signature weights and the output is a natural number that represents the number of mutations the given sample could realistically have. We treat this as a classification problem, where we created 8 different intervals containing different ranges of numbers of mutations from 10 to 10^5^. Then, the neural network is trained using the cross-entropy loss with 8 different classes representing ranges.

SigNet Generator improves upon the current state-of-the art, SynSigGen [27], with its main advantage being that it encodes the main signature correlations existent in the training data (**Extended Data Figure 2**). Instead, SynSigGen independently attributes weight to signatures according to their overall frequencies in the sample set, thus ignoring those correlations. It is important to notice, however, that spurious correlations are created by our method. Future work at the model level will need to be done to address them.

### Data sets

#### PCAWG cancer WGS data

The individual data sets of the PCAWG samples [26] are available at Synapse (synapse.org) and at dcc.icgc.org (https://dcc.icgc.org/releases/PCAWG/). We also used clinical data for each patient, including information about age and sex (https://dcc.icgc.org/releases/PCAWG/clinical_and_histology#!); RNA-Seq data for each tumour (https://www.ebi.ac.uk/gxa/experiments/E-MTAB-5200/); single base substitution signature exposures for each tumour (https://dcc.icgc.org/releases/PCAWG/mutational_signatures). The original PCAWG data set contains 2780 samples from 37 cancer types. For the hypoxia analysis, we used the SigProfiler decompositions of the PCAWG data set and metadata information available in Supplementary Data 1 from Bhandari *et al.* [24]. We also computed hypoxia scores for nine different hypoxic signatures for each of the samples. These results are available in **Supplementary Table 6**. Cancer type abbreviations can be found in Extended Data Table 1 in [26]. For the whole-genome versus whole-exome analysis, we decomposed the mutations that intersect these regions, **Supplementary Table 8**. Similarly for the clonal and subclonal analysis, **Supplementary Table 9**. SigNet’s decompositions for all co-localisation analyses can be found in **Supplementary Tables 11-15**.

#### MC3 cancer WES data

The MC3 mutations [25] for each individual are available at the NCI’s Genomic Data Commons. We used the open-access data files. We also used purity and ploidy information available in TCGA mastercalls.abs tables JSedit.fixed.txt and metadata from Supplementary Table S1, all these available in https://gdc.cancer.gov/about-data/publications/pancanatlas. The MC3 dataset contains 10,186 samples from 33 cancer types. Their decompositions can be found in **Supplementary Table 1**. The hypoxia analysis on the MC3 data was conducted using **Supplementary Table 7**. Cancer type abbreviations can be found at https://gdc.cancer.gov/resources-tcga-users/tcga-code-tables/tcga-study-abbreviations.

#### HMF metastatic cancer data

The HMF database contains whole genome sequencing data from the Hartwig Medical Foundation, which consists of patients with metastatic cancer. This data is available for scientific research upon request and it contains over 4500 patients from more than 38 different primary tumor locations. Their decompositions can be found in **Supplementary Table 2**.

#### Healthy tissue WGS data

This data set contains 561 samples from 28 different tissues and it was extracted from the supplementary materials from [37]. We used Supplementary Table 5 with the whole-genome single base substitutions across all samples, and Supplementary Table 8 with the mutational signatures detected by the authors. SigNet’s decompositions of these samples can be found in **Supplementary Table 3**.

#### GTEx RNA sequencing data

We obtained the mutations called from mRNA sequencing from Supplementary Table 3 in [38]. We also used the raw counts from GTEx Analysis 2017-06-05 v8 RNASeQCv1.1.9 gene reads.gct.gz available at https://www.gtexportal.org/home/datasets to compute the abundances of each trinucleotide context in order to normalise the number of mutations in each category. The number of samples in the GTEx analysis is 253, and they are distributed across 28 different tissues. Their decompositions can be found in **Supplementary Table 4**.

#### Hypoxia scores

In order to use our consensus hypoxia scores we used nine different hypoxic gene expression signatures: Harris, Semenza, Winter, Elvidge, Leonard, Percio, Buffa, Ragnum and Winter2. The first six signatures were obtained from [57] and the other three from [54, 55, 56], respectively. These hypoxic signatures are defined by sets of hypoxic genes containing between 36 and 172 genes, **Supplementary Table 5**.

### Abundance normalisation

The COSMIC mutational signatures were extracted from a mix of whole-exome and whole-genome sequencing data and then rescaled with the trinucleotide occurrences of the whole genome [5]. Since they are therefore not normalised by the number of times each context is present in the human genome, the signatures are specific to the whole-genome context abundances. This means that if we want to analyse a subset of regions of the genome (like in whole-exome or panel sequencing), we need to rescale the number of mutations according to the relative abundances of the whole genome and the specific regions considered. In particular, for whole-exome sequencing data we need to divide the number of mutations in each context by the number of times this context is present in the exome and multiply it by the number of times it is present in the genome. SigNet Refitter has this already implemented, such that the user can specify if their input data correspond to the whole exome or the whole genome and the algorithm will rescale the data accordingly.

If the analysis involves another subset of the genome, we need to specify what are the abundances that should be used. This means that we need to input the number of times each trinucleotide context is present in the region we are calling the mutations from. For the GTEx analysis, this meant that we needed to compute the abundances of the trinucleotide contexts for the genes expressed in each tissue from which the mutations were extracted. In order to do so, we used the raw gene expression counts from the RNA-seq data and computed the mean number of reads for each gene across all the samples in the tissue. With this, we obtained an estimation of how expressed each gene is in each tissue. We used all genes with expression larger than 100 counts in order to ensure mutations could be detected. To obtain this threshold we considered that the median variant frequency is 0.05 and at least 4 counts from the variant allele need to be sequenced in order to call the mutation [38]. Then, for each tissue, we computed its total abundances by summing each trinucleotide abundance for all genes that exceeded the threshold.

For the co-localisation analyses, we had to compute the abundances of the regions considered within each group. For example, in the replication time analysis, we had to determine the frequencies of all trinucleotide contexts per set of genomic regions contained in each of the replication time bins (ranging from early to late). This is a crucial step that we repeated in all the different analyses that focused on mutations from specific regions, i.e., that did not take into account the mutations from the entire genome.

### Mutational signatures and clonality

In order to identify the mutational processes in early and late development in cancer, we can look at clonal and subclonal mutations, respectively. In this analysis, we used the mutational data from PCAWG [26]. For a diploid site, subclonal mutations can be identified by using a test with the following assumptions:

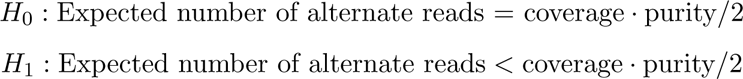

This hypothesis can be assessed by conducting a binomial test with parameters *n* = coverage and *p* = purity*/*2. If the null hypothesis (*H*_0_) cannot be rejected, the mutation is considered clonal. Otherwise, it is categorized as subclonal. The factor of 1/2 within the probability *p* of the binomial test comes from assuming that the cells are diploid. This is a fair assumption in healthy cells, but it is not generally true in cancer. For this matter, we first needed to filter out the mutations that were in regions where the copy number is different from 2. Since we are removing regions from the genome, we needed to recompute the abundances for each sample and correct the mutation counts before applying the signature decomposition. Moreover, given that the subclonal group typically contains fewer mutations than the clonal group, we downsampled the mutations in the clonal group to match the subclonal group’s numbers.

Then, we ran SigNet Refitter separately for each class. This yielded a signature decomposition for each sample and class (one for clonal mutations and another one for subclonal mutations). We excluded samples classified as “unfamiliar” (which would have otherwise been fit with NNLS). After the filtering, we had 2323 samples in the clonal and 1927 in the subclonal group. We further restricted the samples to those that were paired, so those that had a realistic decomposition in the clonal and in the subclonal part, which resulted in 1767 samples. In order to assess statistical significance of the results we used the Brunner-Munzel test, excluding paired samples with 0-0 values, since in such samples the mutational signature is not active.

### Differences between whole-exome and whole-genome mutational signature activities

To explore the prevalence of different signatures within regions of the genome, we conducted a comparative analysis of signature decomposition among the entire genome, genes, and exons. To generate the last two groups, we used bedtools to intersect the mutations in whole-genome samples with the coordinates of the protein-coding genes and exons, respectively. To ensure comparability between the three groups, we balanced the mutation counts in the entire genome and genes by downsampling them to match the number of mutations in exons. Then, we ran SigNet Refitter for all samples in the three groups while considering their respective trinucleotide abundances. Additionally, we transformed the decomposition results by multiplying them by the total number of mutations, converting the weights into mutation counts. Once again, we filtered out samples classified as out-of-training-distribution (i.e., “unfamiliar”) by SigNet Detector. Finally, we restricted our analysis to samples with reliable information across the three groups, resulting in a dataset comprising 1354 samples.

We compared these results by computing the mean mutation count by mutational signature and by cancer type. Since we are comparing whole-genome with whole-gene and whole-exome data, we rescaled the total number of mutations so that it is comparable across sets. To do so, we divided the number of mutations in the PCAWG data set by the whole genome size divided by 10^7^, and similarly for the other two groups, dividing by their region size divided by 10^7^ (**Figure 4**). We again computed statistical significance using the Brunner-Munzel test, excluding paired samples with 0-0 values, since in such samples the mutational signature is not active.

### Hypoxia Analysis

The analysis of the correlations between the mutational signatures and hypoxia was conducted similarly to Bhandari *et al.* [24], with three changes: we used the total number of mutations attributed to each signature (i.e., the activity) instead of weights, we included the coverage as an explanatory variable of the model, and we used a consensus hypoxia score. It is important to use the total number of mutations instead of signature weights because weights are compositional data. This means that any absolute increase in one signature must produce a decrease in the relative weights of all the other signatures even if their total contributions did not change. Using a consensus hypoxia score is important because there is not yet a validated set of hypoxic genes. Bhandari *et al.* used the Buffa hypoxia score [54] in their analysis. However, there exist many different scores that provide a set of hypoxia-associated genes, and they contain different genes with very limited overlap. The most well-known hypoxia scores are Buffa, [54], Ragnum [55] and Winter [56], but other sets are also used [57]. Here we used a consensus approach to determine which mutational signatures are associated with hypoxia: Similarly to what is done with mutation callers, we say that a signature is significantly correlated with hypoxia if it is significantly correlated with at least two different hypoxia scores. In total, we used nine different hypoxia scores. The three aforementioned, and also Harris, Semenza, Winter (a second one), Elvidge, Leonard and Percio [57]. In order to compute the hypoxia scores from the set of hypoxia-associated genes, we ranked gene expression data in the same way as described in Bhandari *et al.* [24]. Specifically, for each gene in a hypoxia signature, we assigned a value of +1 to the patients with the top 50% of mRNA, and a value of -1 to those in the bottom 50%. Then, we assigned a hypoxia score for each patient by adding all the +1 and -1 for all genes in the given hypoxia signature.

As in Bhandari *et al.*, we used a linear mixed-effects model that associates hypoxia with each of the different mutational signatures, while taking into account other covariates, including tumour purity, patient age, patient sex and the average tumour sequencing coverage. Together with the purity, the latter can explain the technical ability to call mutations. In addition, the cancer type was included as a random effect in the model. We then compared each of the full models to a null model that did not contain the mutational signature as an explanatory variable. Using an ANOVA, we can determine whether the given mutational signature has a significant effect on describing hypoxia and, therefore, whether there is an association between its number of mutations and hypoxia. More concretely, we compared the full model for each signature SBSx:

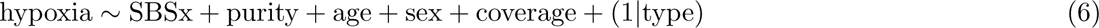

to a null model without the signature:

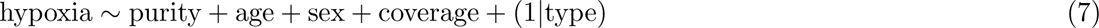

Then, we computed the p-value for each of the comparisons and corrected them by multiple testing using FDR adjustment. In **Figure 3**, we show the signatures that had an adjusted p-value lower than 0.05 after the correction.

For the MC3 data set, we conducted the same analysis as described for PCAWG, but here we used SigNet Refitter for the signature decomposition first. We again filtered out those samples that were classified as “unfamiliar” by SigNet Detector to avoid estimation of weights with NNLS only. After filtering out those samples and keeping samples for which we could gather information for all the covariates in our model, 7411 samples remained in the analysis in **Figure 3b**.

We also conducted the same analysis as in [24] by only exchanging the weights for total numbers of mutations (i.e., keeping the Buffa score and not using the coverage as one of the explanatory variables). These results are shown in **Extended Data Figure 9**.

### Mutational signatures present in different parts of the genome

We conducted several analyses to investigate the co-localisation of certain signatures with specific genetic and epigenetic features. In particular, we analysed the mutational signatures present in regions characterised by early and late replication, regions with varying levels of insertions, deletions or structural variants, and regions where different histone modifications occur. All of these analyses employed a very similar methodology.

First, we identified the genomic coordinates of the different groups within the specific analysis. For instance, in the case of histone modifications, we retrieved the ChIP-seq peak coordinates that would define the regions under study, with the remainder of the genome constituting the comparison group. Second, we intersected the coordinates from the different groups defined in the initial step with the mutations observed in the PCAWG samples. Third, we constructed the mutation count matrix by collapsing the trinucleotide context of each mutation into the pyrimidines, counting the occurrence of each type of mutation within each cancer type. In other words, we aggregated all mutations from the same cancer type across all samples to generate a mutation count data frame where each row corresponds to a cancer type rather than an individual sample. We undertook this approach due to the small size of the genomic regions in at least one of the groups in various analyses, resulting in either zero or very few mutations per sample that intersected with these regions. Fourth, in instances where different groups exhibited substantially different numbers of mutations, we downsampled the mutations from the larger datasets to match the count of the smallest one. This step was taken to mitigate the challenges associated with variations in accuracy during the signature decompositions, which could be influenced by substantial differences in noise levels across the samples. Fifth, we normalised the mutation counts for each cancer type based on the abundances of the regions under study, as previously described.

These five steps are essential for generating the appropriate input for our signature decompositions. With this, we can execute SigNet and subsequently compare the prevalence of different mutational signatures among the various groups. To do this, we first converted the signature weights obtained from the algorithm into total mutation counts. This step is essential for eliminating the compositional nature of the weights, which could otherwise lead to spurious results. To achieve this, we multiplied each signature’s weight by the original total number of mutations within the cancer type’s genetic coordinates for the given group. We then divided this product by the total length of the region, ensuring that the mutation counts remain comparable across regions of varying lengths. Then, we scaled the values in each bin per each cancer type by the mean value of the cancer type across bins to make the different cancer types comparable, before computing the mean across cancer types per bin (**Figure 5**). Error bars were computed as the standard error of the mean. In the following sections, we provide detailed information on the specific data and techniques employed in each of our analyses.

#### Replication time

We used the ENCODE Repli-seq wavelet-smoothed signal data, which we downloaded from the UCSC portal (https://genome.ucsc.edu/cgi-bin/hgTrackUi?db=hg19&g=wgEncodeUwRepliSeq). Consistent with prior studies, [104], we averaged the results from the following cell lines: GM12878, HeLa, HUVEC, K562, MCF-7 and HepG2. In this dataset, replication peaks correspond to replication initiation zones (early replication), and valleys correspond to termination zones (late replication). For clarity in the main figures, we show the results from early to late replication times.

We also excluded all zero and low values from the data, as they appeared to be artefacts or missing data. Finally, we divided the data into five groups of equal replication time value range sizes.

#### Histone marks

We leveraged all the high-quality ChIP-seq data available in the ENCODE Portal (https://www.encodeproject.org/). This includes data from eight different laboratories and 31 distinct histone modifications obtained from various cell lines. We retrieved the peaks corresponding to each histone modification from the ENCODE Portal and conducted a liftover to convert their coordinates from GRCh38 to GRCh37, to ensure compatibility with the mutation data from the PCAWG.

Since the peaks of different histone modifications are located within regions characterized by varying replication time values, and considering the well-established strong correlation between replication time and mutability, we accounted for this effect by adjusting the total number of mutations within and outside the histone peaks based on the expected number of mutations corresponding to the given average replication time value. To achieve this, we initially computed a linear regression analysis for each cancer type. We utilized the mutation counts per signature within each replication time bin, as obtained from the replication time section of our analysis. This process allowed us to establish a linear function capable of predicting the number of mutations for a specific signature within a particular cancer type, given a replication time value, **Supplementary Table 10**. Using the average replication time as well as the whole-genome average, we made predictions for the number of mutations both within and outside the histone regions. These predictions were then used to correct the total mutation counts accordingly.

Lastly, we performed a Wilcoxon rank-sum test to assess the statistical significance of differences between the two groups for each signature and implemented FDR multiple testing correction.

#### Indels and structural variants

We retrieved the data from the ICGC data portal, acquiring both ICGC and TCGA samples’ data (https://dcc.icgc.org/releases/PCAWG/consensus_snv_indel, https://dcc.icgc.org/releases/PCAWG/consensus_sv). For the indels analyses we used the PCAWG maf files that contain single nucleotide variants, multiple nucleotide variants and insertions and deletions. From these, we kept only the insertions and deletions of size 1, and the deletions of 5 base pairs or more. For the structural variants analyses we filtered out the deletions to avoid biased results driven by different target sizes between donors with and without the deletion, since they were very big and sometimes spanning several windows.

We divided the genome up into non-overlapping windows of size 500 kbp and defined two groups: windows without any variant events of interest (group 0), and windows containing 1 or more such events (group 1), at the sample level. Additionally, for each cancer type, we restricted to the windows that had at least one event of the specific variant in at least one of the samples. Subsequently, for each focal signature, we selected only the samples with weight larger than zero in the whole-genome decompositions derived from SigNet. For every genomic window, we then matched the number of samples in the two groups by selecting, randomly, samples from group 0 to match the number of samples in group 1 at the given genomic location, thus controlling for correlation of indel rates with mutation rate covariates. Then, we downsampled the mutations of each sample such that the activity of the focal signature corresponds to the minimum activity of that signature across all samples from the cancer type. This step was necessary to avoid biases from pooling mutations across samples with different activities. Next, we pooled all the SNPs from all samples in the same cancer type and all windows in the same group (0 or 1) and we downsampled a second time, the mutations from the biggest group (usually group 1) to match the number of mutations in the smallest group (usually group 0). The final steps encompassed abundance normalisation and SigNet decomposition, as previously described.

The quantity shown in **Figure 5b** corresponds to a scaled average multinomial success probability of the signature activity in windows with and without an event. For each event category (0 and 1), we compute this metric, *q*, by dividing the total mutation count attributed to the focal signature by the whole-genome activity of the signature per sample (which is the same for all samples due to the downsampling described earlier) multiplied by the total number of windows across all samples that were considered in the given cancer type. Furthermore, since we consider different genomic windows for each of the cancer types, we scale the results of each cancer type by the average *q* of the given cancer type between group 0 and group 1. This ensures a fair comparison across different regions of the genome, preventing results from being skewed towards more mutable regions.

To quantify uncertainty, we employed a bootstrap resampling approach at the cancer type level, by bootstrapping 100 times the mutations per cancer type. Since cancer types with very few mutations in the selected windows have very noisy decompositions, we computed the average across cancer types by weighting the results by the inverse of their variances, where the variance was determined through the bootstrap process. This means that the height of the bars and the error bars in **Figure 5b** are not the regular mean and standard error (SE), but rather the inverse-variance-weighted mean and standard error, which correspond to

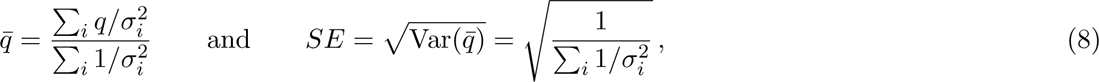

where the sum runs over all cancer types *i*. Cancer types that were classified as “unfamiliar” by SigNet in more than 10% of bootstrap replicates were removed from the computation of the weighted mean because they had unreliable estimation of the variance. For those cancer types that had a variance equal to zero, we updated the value to the minimum value of the variance across all cancer types that had a variance larger than zero, to make the computation numerically possible.

To assess significance, we used a weighted t-test applied to the difference between the weighted means for each variant type. For this, we defined the t-statistic as

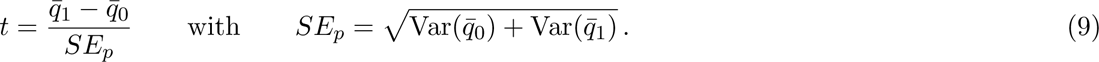

### Mutational signatures and POL*θ*

To investigate the association between distinct mutational signatures and POL*θ*, we conducted an analysis aimed at quantifying the correlation between POLQ gene expression and the load of these signatures. First, we applied quantile normalisation to the Transcripts Per Million (TPM) values obtained from the MC3 gene expression dataset. This normalisation procedure is a standard practice in the context of gene expression analysis, as it ensures the comparability of values across genes and samples. Following normalisation, we established pairs of TPM values for the POLQ gene and the total mutation counts associated with SBS3, SBS5, and SBS40 for each sample. Subsequently, we computed the Pearson correlation coefficients for these pairs, revealing highly significant correlations for all three mutational signatures. To further refine our findings, we employed a correction method for the p-values obtained from these correlation tests. To do so, we compared the correlation coefficients to a null distribution of correlation coefficients derived from all genes’ expression data. This null distribution was computed separately for each of the three mutational signatures. Subsequently, we calculated a corrected p-value by determining the proportion of genes with correlation coefficients greater than the one observed for the POLQ gene. These corrected p-values are presented in **Extended Data Figure 10**.

### Curveball

We here provide an intuitive derivation of the inferred relationship between mutation status of a gene and signature activity under the curveball algorithm for a system of three genes (1, 2 and 3) and two samples (A and B), with the contingency matrix

|  |  |  |
| --- | --- | --- |
| $m_{1A}$ | $m_{1B}$ | $g_1$ |
| $m_{2A}$ | $m_{2B}$ | $g_2$ |
| $m_{3A}$ | $m_{3B}$ | $g_3$ |
| $s_A$ | $s_B$ | |

where *m_ij_ ∈ {*0, 1*}* denotes the mutation status of gene *i* in sample *j*, while *g_x_* and *s_x_* are the row and column total, respectively. Since we want to test the outcome in case the mutation probabilities vary across genes and samples, let us assume that gene 1 and 2 have the same basal mutation rate, but gene 3 is more mutable. Further, sample B has a higher mutation rate than sample A. However, we shall assume no explicit dependence of the mutation load of a sample on the mutation status of either of the genes, i.e., the mutation probabilities of the three genes are independent. Then, we can define the mutation probability of gene *i* in sample *j* as follows:

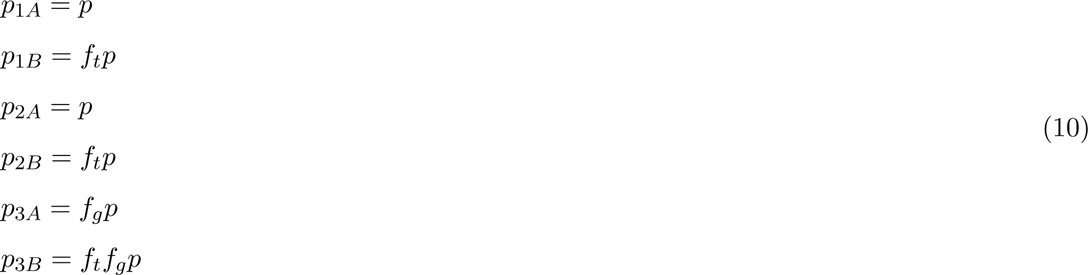

where *p ≪* 1, *f_t_* ≲ *O*(10) is a tumour-specific factor multiplying the basal mutation rate *p* for all genes in sample B and *f_g_* ≲ *O*(10) is a gene-specific factor multiplying the basal mutation rate *p* for gene 3 in all samples.

For a system of three genes and two samples, there exist 64 possible configurations of the contingency matrix. The curveball algorithm randomises blocks of structure (0,1)(1,0) or (1,0)(0,1), thus preserving row and column totals. Out of the 64 possible configurations, 18 have blocks which can be randomised. For each gene, out of these 18 configurations, only six produce a difference between the observed and expected mutation load as a function of gene mutation status. It can then be shown that the events in which the observed load is positively associated with the mutation status of gene 3, producing a larger relationship than expected, are on average more likely than the events in which the relationship is more negative than expected. The inverse holds for genes 1 and 2. In words, the effect of a mutation on gene 1 or 2 on the mutation load of the sample would be underestimated, while the effect of a mutation on gene 3 on the mutation load of the sample would be overestimated. **Extended Data Figure 17** shows simulation results for a realistic number of genes and tumors.

## Code availability

The code we developed for all three SigNet modules along with installation guidelines and an extensive documentation is available at https://github.com/weghornlab/SigNet. There are different ways of installing SigNet, but we recommend using a Singularity image that we created with the necessary packages to execute the algorithm. This file, *signet.sif*, can be downloaded from https://weghornlab.net/Software/signet.sif. We recommend using Singularity3.1. The easiest way to install and run SigNet is by executing:

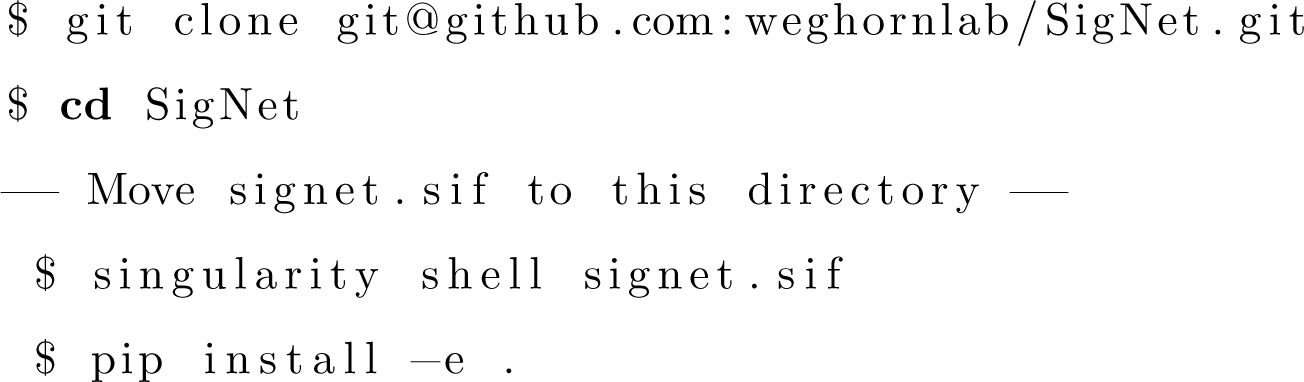

Afterwards one can check that the installation worked properly by running:

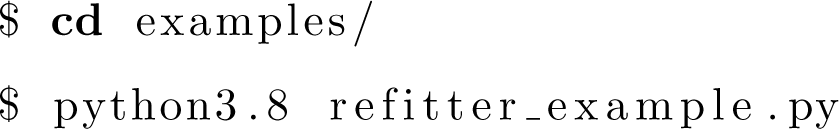

## Acknowledgments

We acknowledge support of the Spanish Ministry of Science and Innovation through the Centro de Excelencia Severo Ochoa (CEX2020-001049-S, MCIN/AEI /10.13039/501100011033), and the Generalitat de Catalunya through the CERCA programme. This work was also funded by the Spanish Ministry of Science and Innovation through grants PGC2018-100941-A-I00 and PID2021-128976NB-I00. We thank Shamil Sunyaev for a critical reading of the manuscript as well as all members of the Weghorn Lab for helpful discussions and suggestions. This publication and the underlying study have been made possible partly based on data that Hartwig Medical Foundation and the Center of Personalised Cancer Treatment (CPCT) have made available to the study through the Hartwig Medical Database.

## Conflicts of interest

The authors declare no competing interests.

## Extended Data

**Extended Data Figure 1:**
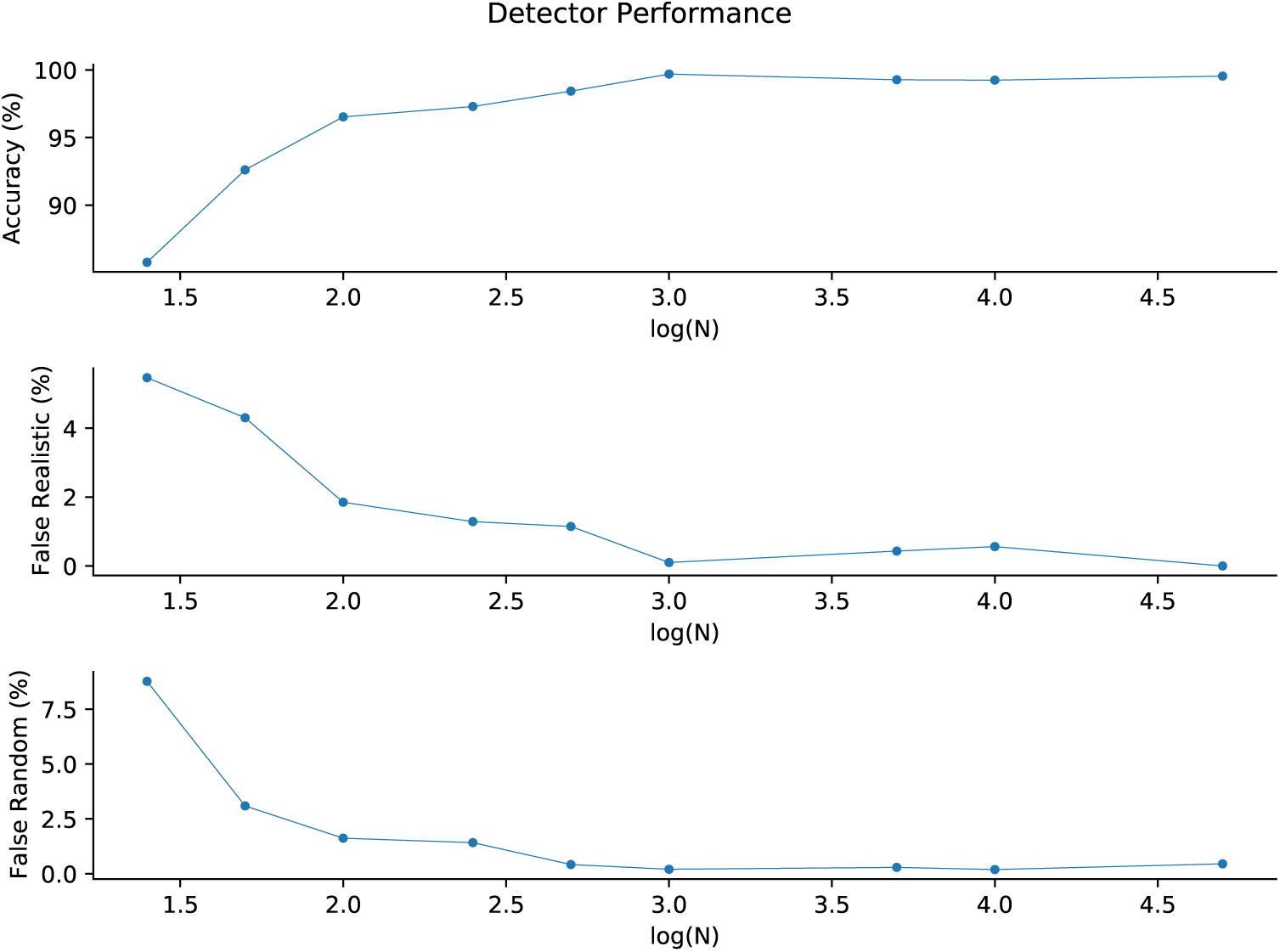
Detector performance versus the number of mutations. The accuracy of the detector increases with the numbers of mutations because the samples become less noisy. Similarly, the number of samples (out of all samples) falsely classified as random and as realistic decreases with the number of mutations. The number of mutations, *N*, is provided in logarithmic scale.

**Extended Data Figure 2:**
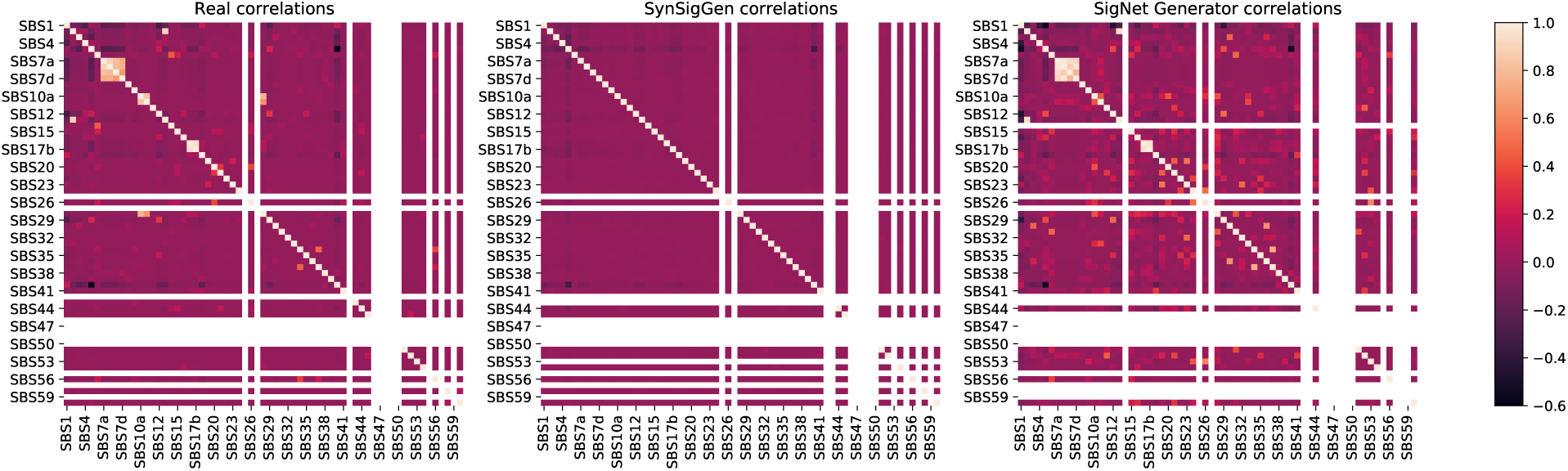
Pairwise signature correlations in real and synthetic data. Pearson correlation matrices for PCAWG signature assignments [26] (left), SynSigGen [27] synthetically generated signature combinations (centre) and SigNet Generator output (right). Notice how SynSigGen misses all signature correlations, while SigNet Generator maintains the most prominent correlations. At the same time, SigNet Generator exaggerates certain correlations.

**Extended Data Figure 3:**
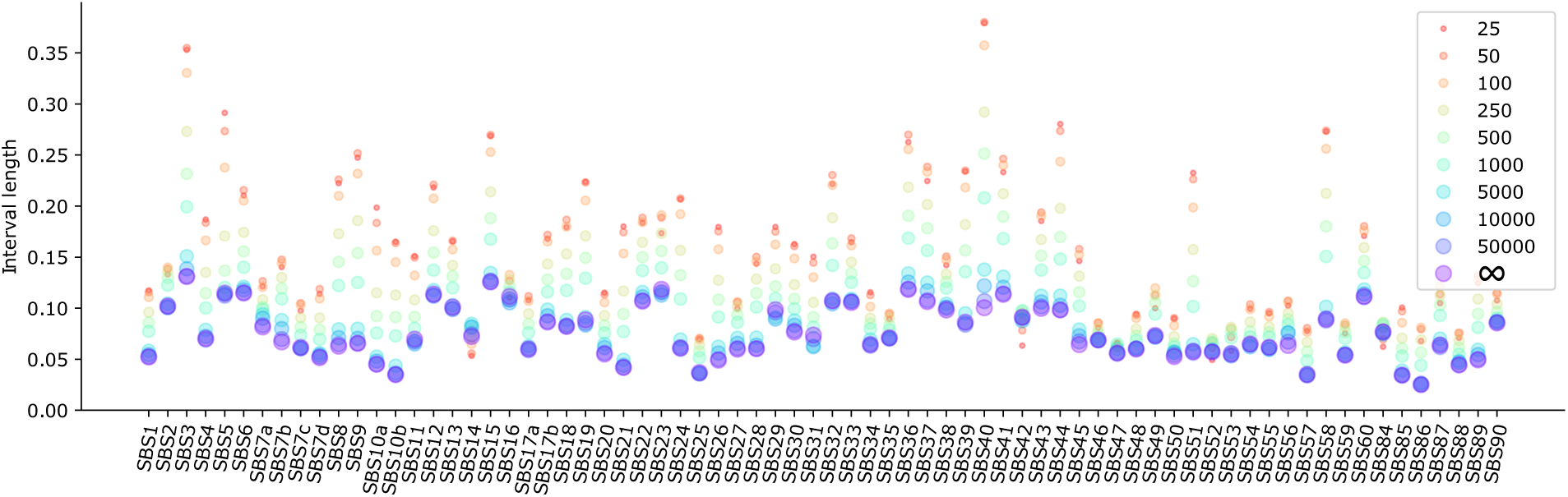
Interval width of the error intervals generated by the ErrorFinder module in SigNet Refitter. Shown as a function of the mutational signature and the number of mutations of the sample. The width of the interval decreases with larger number of mutations because the samples are less noisy, and therefore, easier to decompose. Signatures like SBS3, SBS5 and SBS40 present larger intervals because of their flat shape, which makes them more difficult to identify than the rest.

**Extended Data Figure 4:**
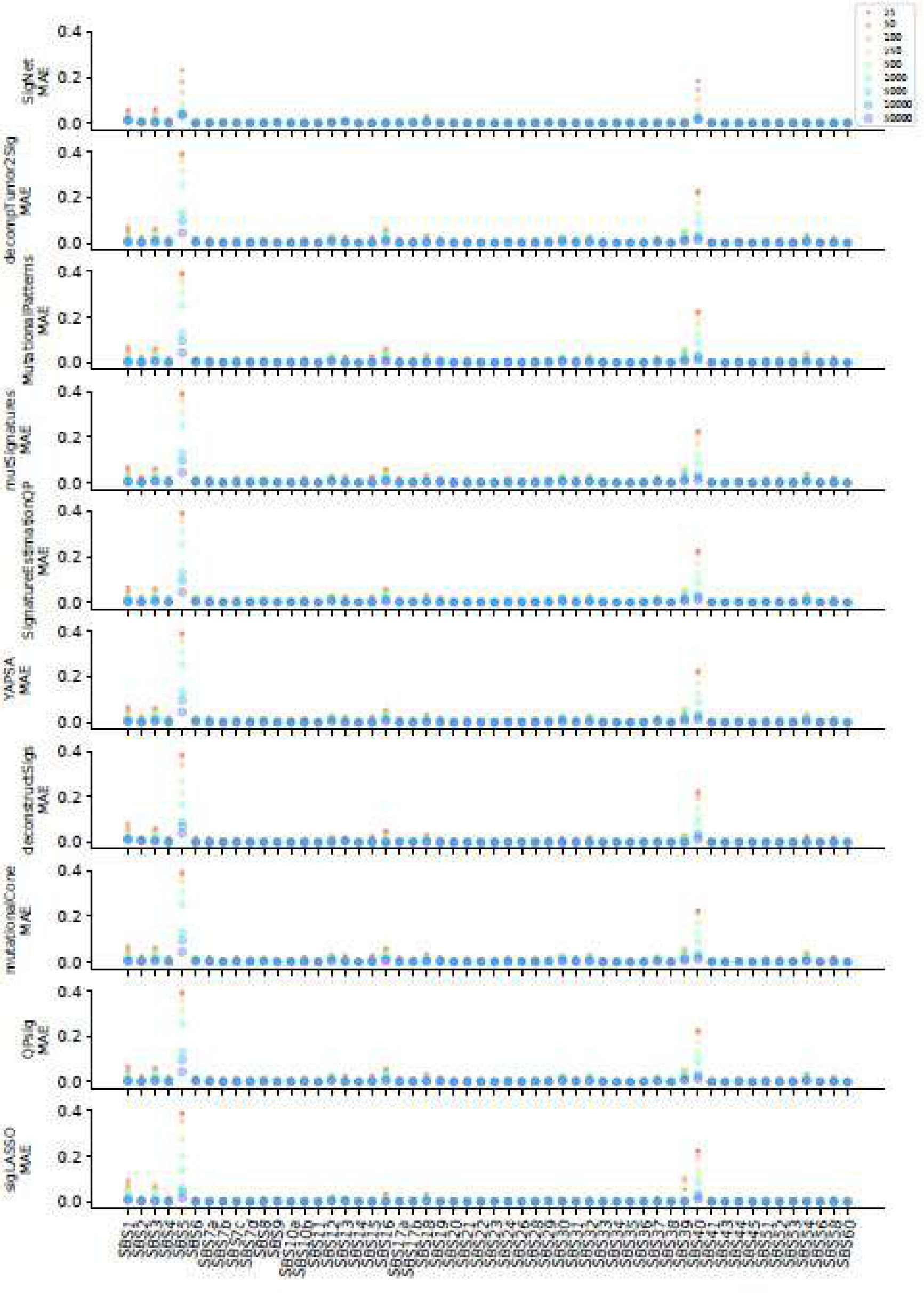
Mean absolute error (MAE) between the weight guess and the label. Shown as a function of the mutational signature and the number of mutations in the sample for all the tested methods. The top plot (SigNet) is the same as Figure 1f.

**Extended Data Figure 5:**
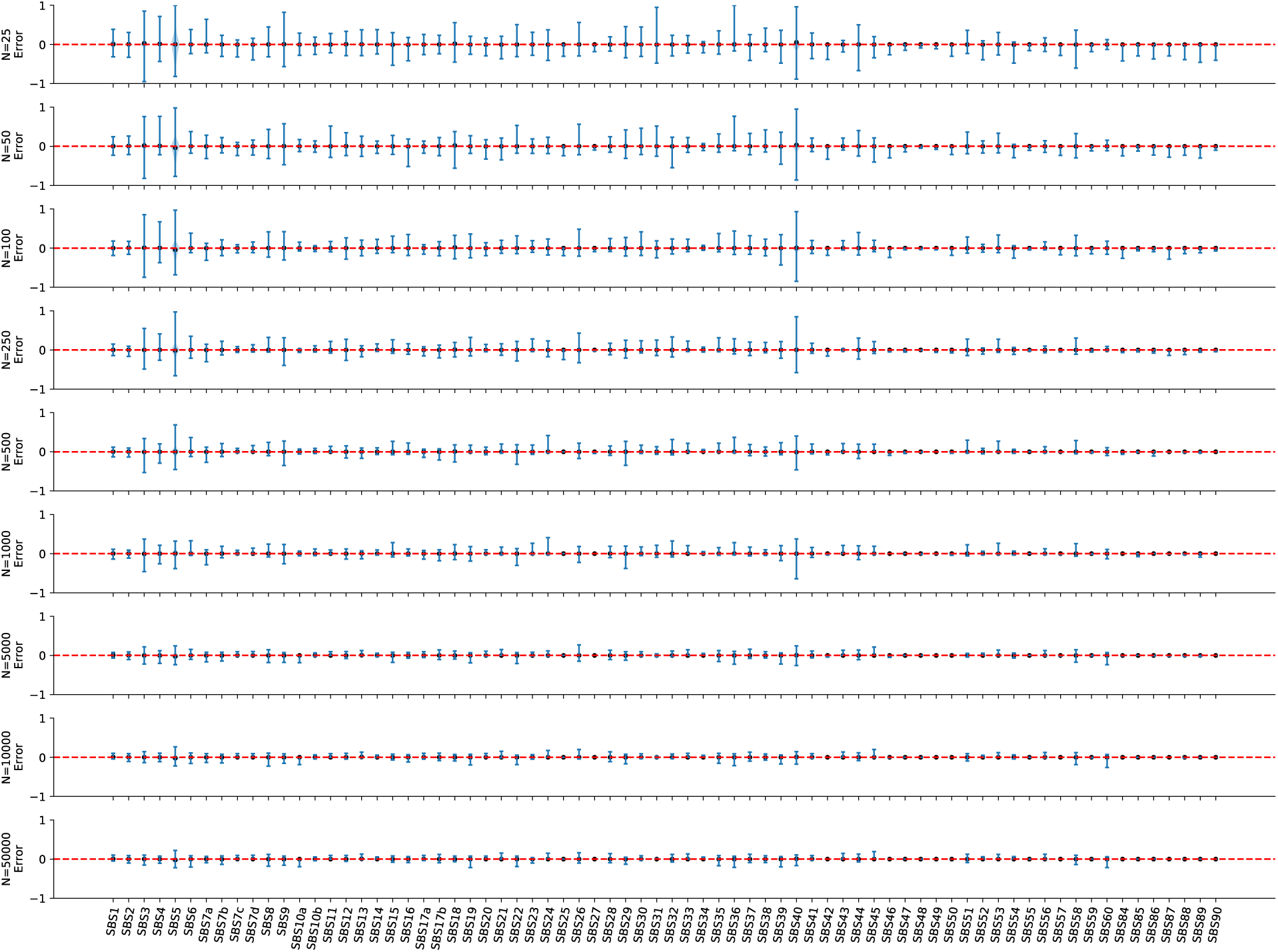
Violin plots showing the distribution of the error between the labels and the guesses of the test set. Shown for each mutational signature and separated between different numbers of mutations (N=25 to N=50000 in rows in increasing order). The red line denotes zero error and the black dots the mean error for each mutational signature.

**Extended Data Figure 6:**
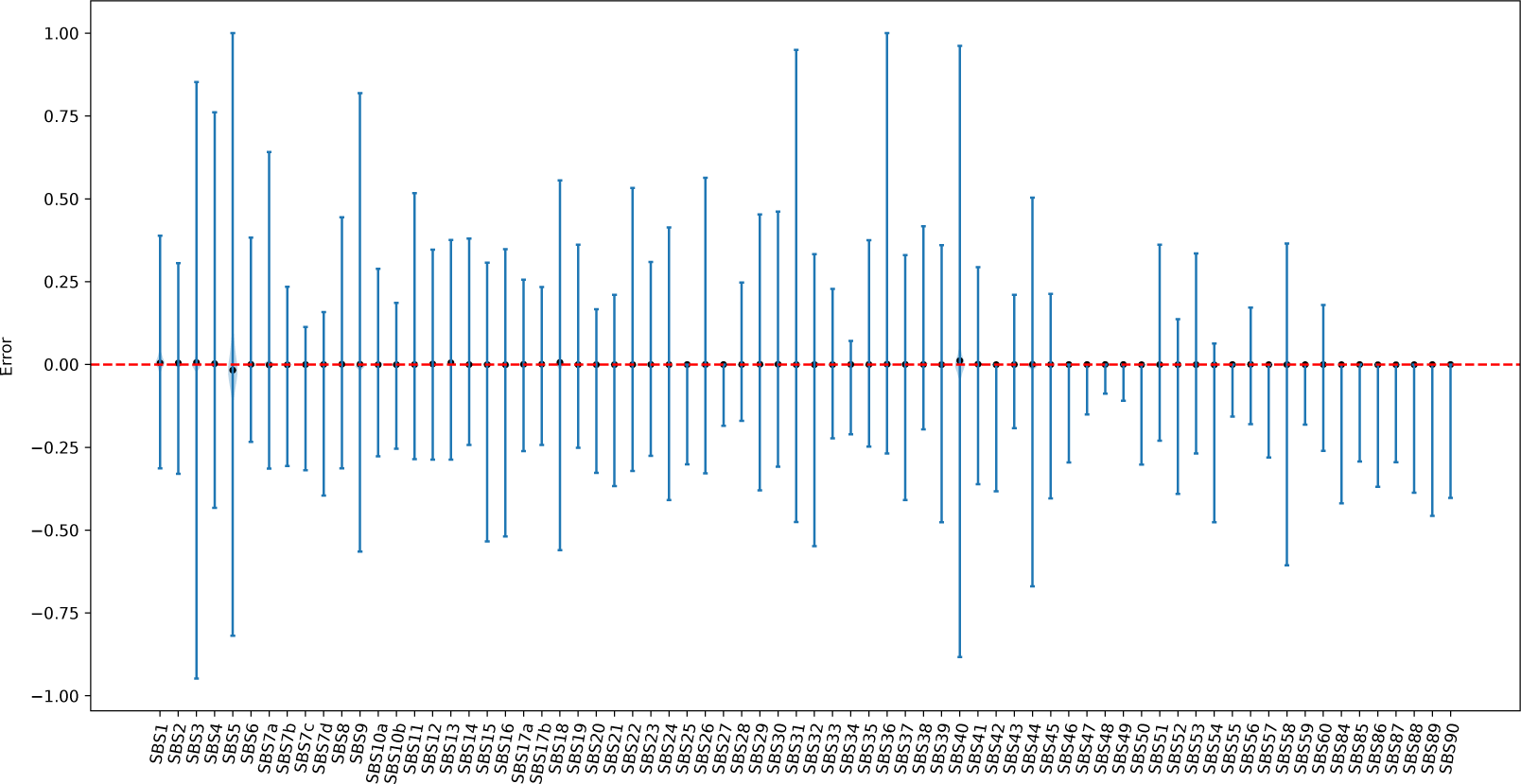
Violin plot showing the distribution of the error between the labels and the guesses of the test set. Mean for each mutational signature across all *N*. The red line denotes zero error and the black dots the mean error for each mutational signature.

**Extended Data Figure 7:**
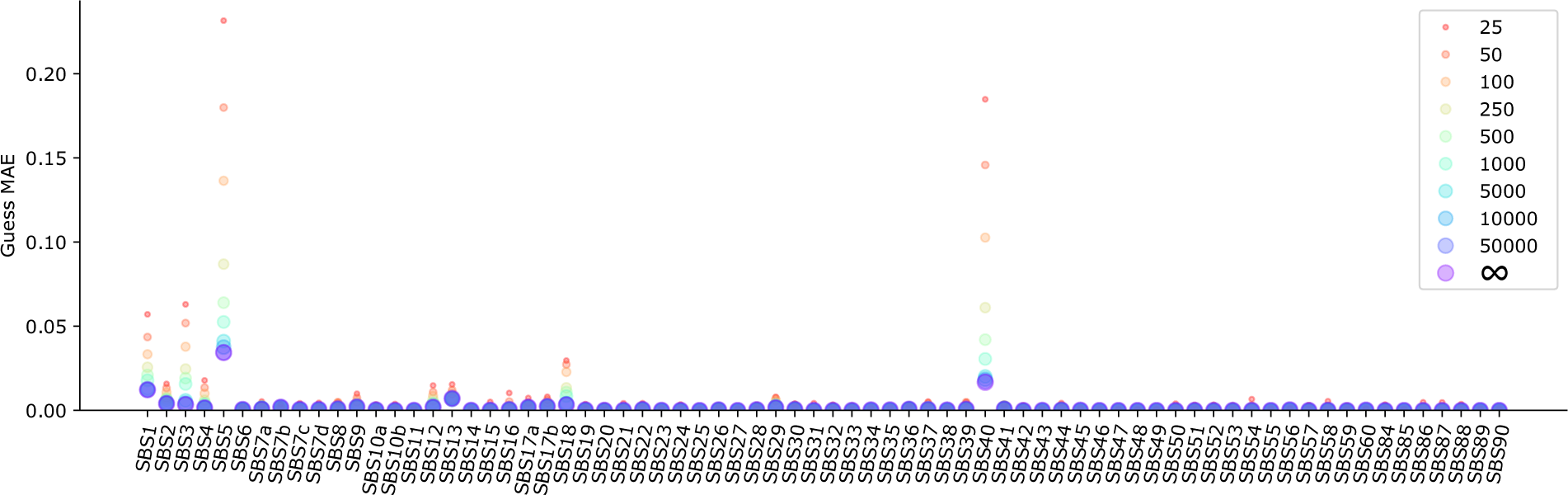
Mean absolute error (MAE) between the guess and the label for all the mutational signatures available in COSMIC v3.1. Shown as a function of the number of mutations in the sample. The error of the guesses decreases with larger number of mutations. Signatures like SBS5 and SBS40 have larger errors because of their flat shape that makes them more difficult to identify than the rest. Note that the signatures that are not present in any of the PCAWG training samples will be learnt as never present. Therefore, the Finetuner part of the algorithm will not include them.

**Extended Data Figure 8:**
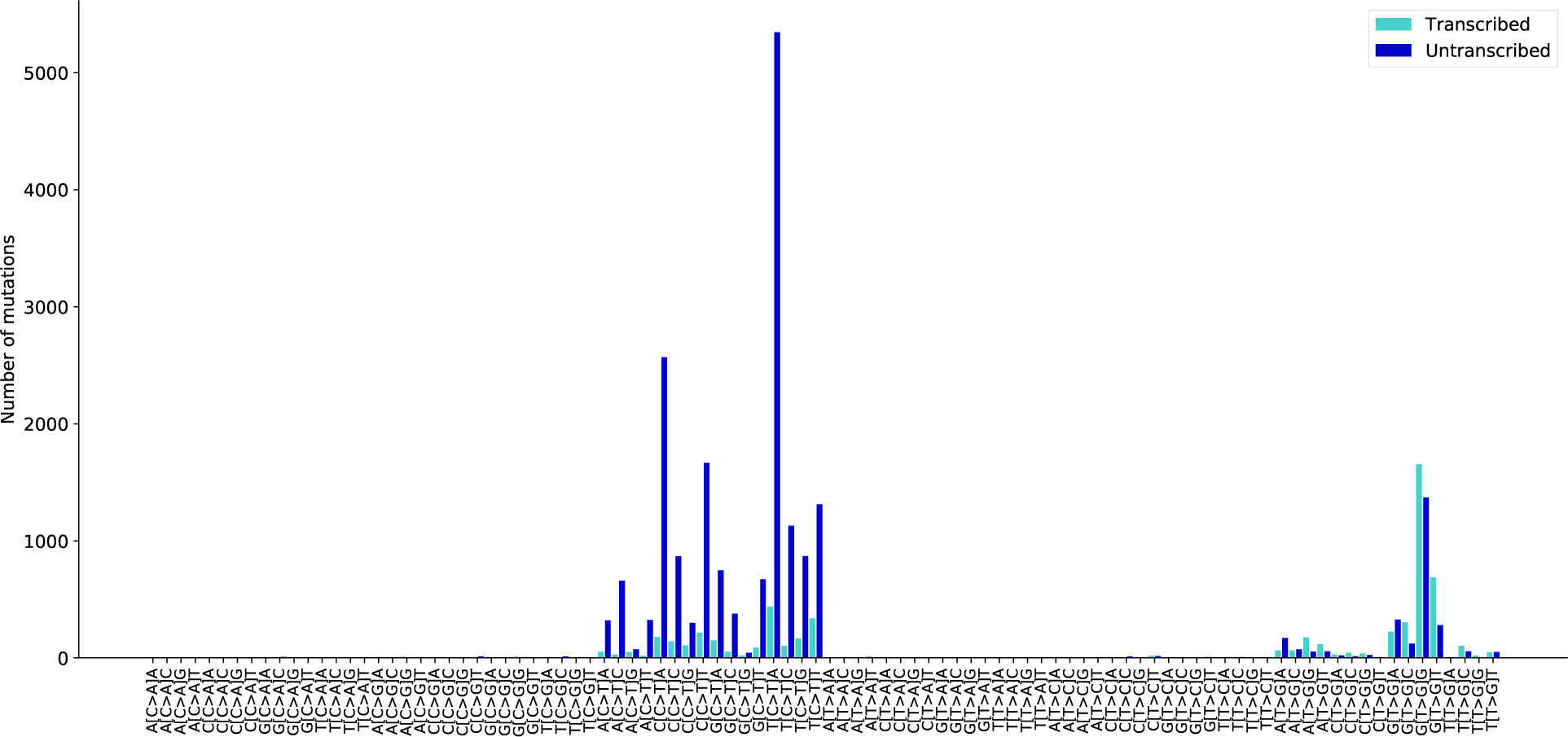
Transcription strand bias in unusual mutational signatures observed in RNA-seq data. Pooled number of mutations attributed to SBS2, SBS7a, SBS7b, SBS19, SBS30 and SBS60 over 100 independent assignments. The assignments were conducted for all mutations in each of the 14 tissues from the GTEx data set that were classified as realistic and they were polarised by transcription strand.

**Extended Data Figure 9:**
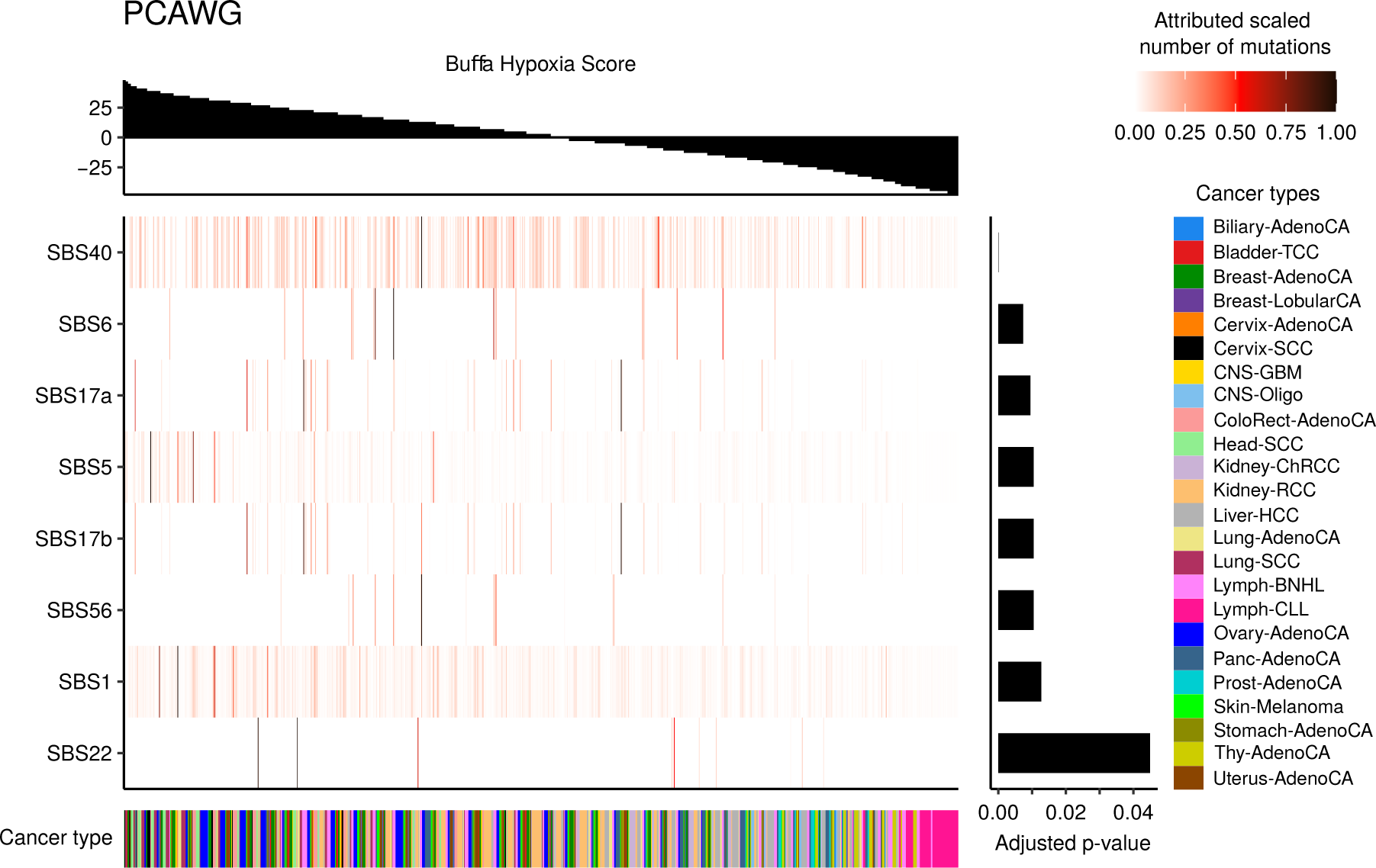
Association between hypoxia and signature mutation count (activity) instead of weight. The total number of mutations attributed to each mutational signature is obtained from the SigProfiler assignments in the PCAWG data [26], measured and sorted by FDR-adjusted p-value. Hypoxia is measured with the Buffa score and significance is based on an ANOVA test, as in [24].

**Extended Data Figure 10:**
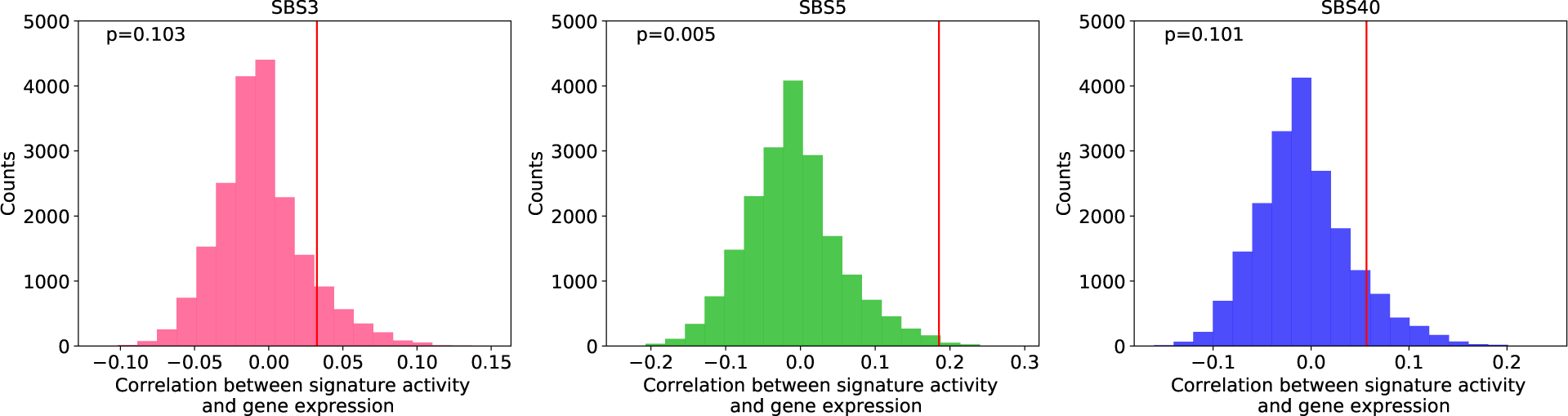
Histograms illustrating the Pearson correlation coefficients between quantilenormalized TPM values of all genes and the mutation load associated with SBS3, SBS5, and SBS40 (left to right). The vertical line represents the correlation coefficient value for the POLQ gene, and the p-value displayed in the upper left corner of each plot signifies the proportion of genes exhibiting correlation coefficients greater than that of the POLQ gene.

**Extended Data Figure 11:**
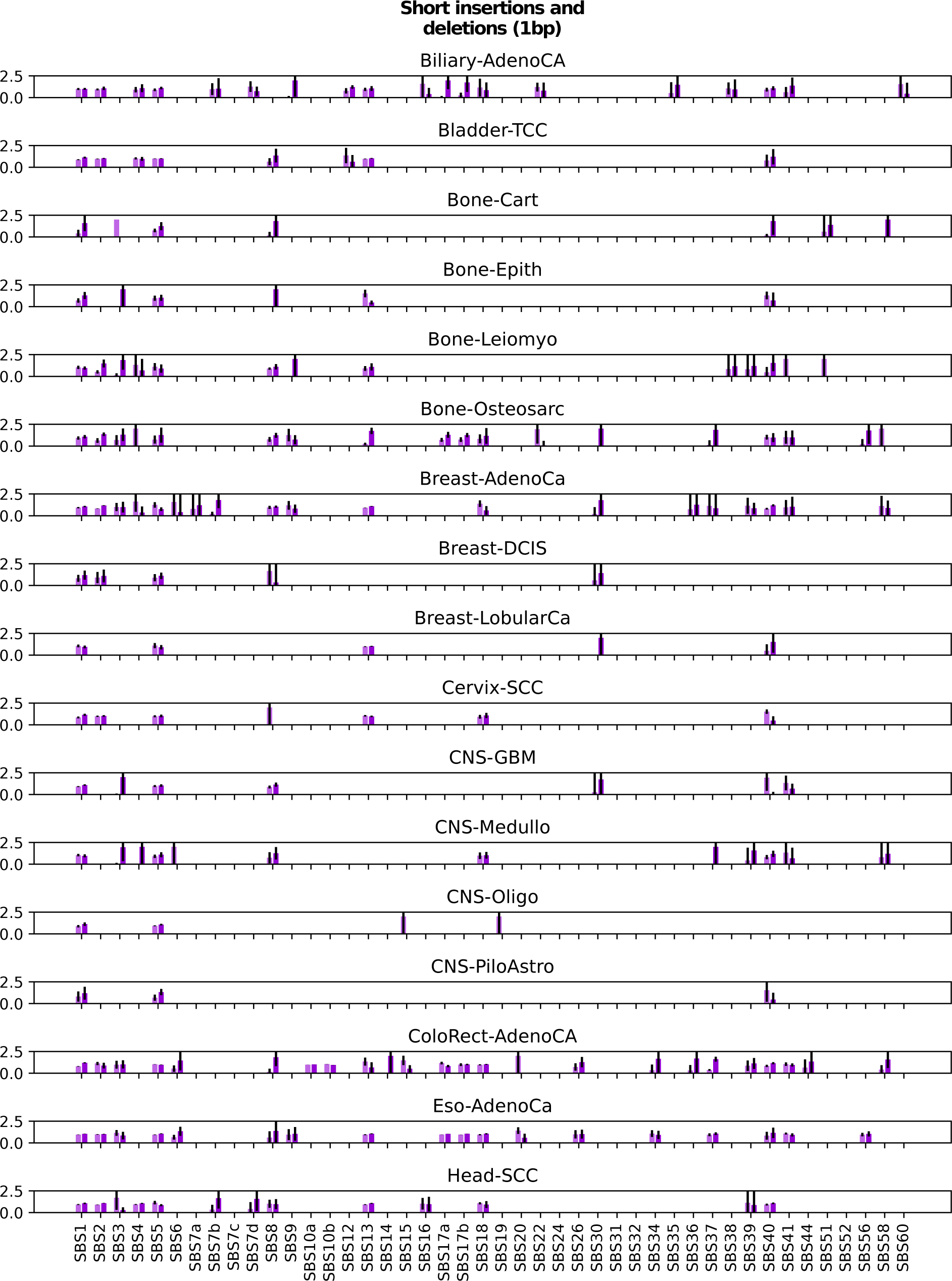
Scaled mutability of windows without short indels (lighter color) and windows with short indels (darker color) for all signatures for the first half of all cancer types in PCAWG. Error bars are computed through 100 bootstrap replicates.

**Extended Data Figure 12:**
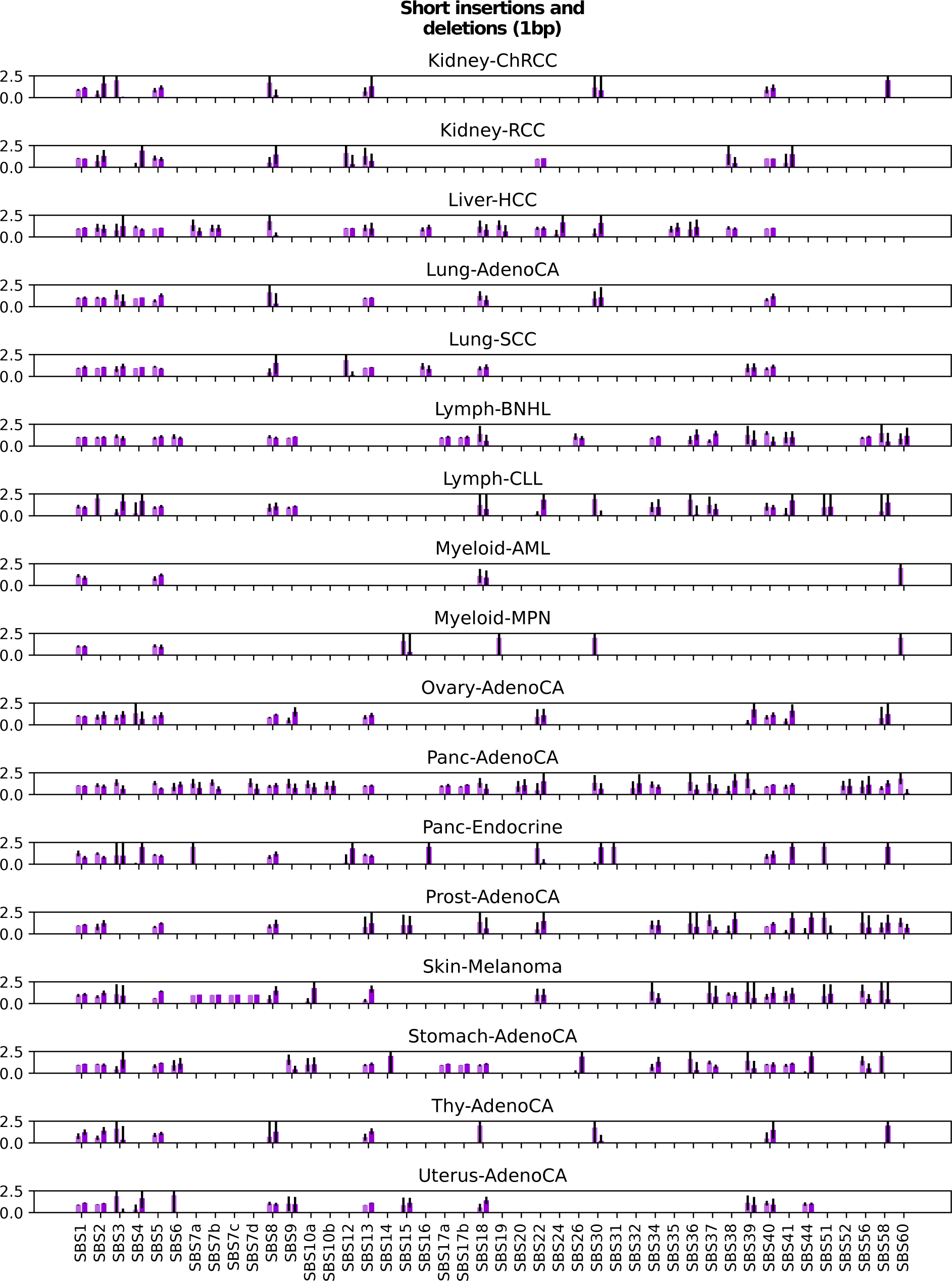
Scaled mutability of windows without short indels (lighter color) and windows with short indels (darker color) for all signatures for the second half of all cancer types in PCAWG. Error bars are computed through 100 bootstrap replicates.

**Extended Data Figure 13:**
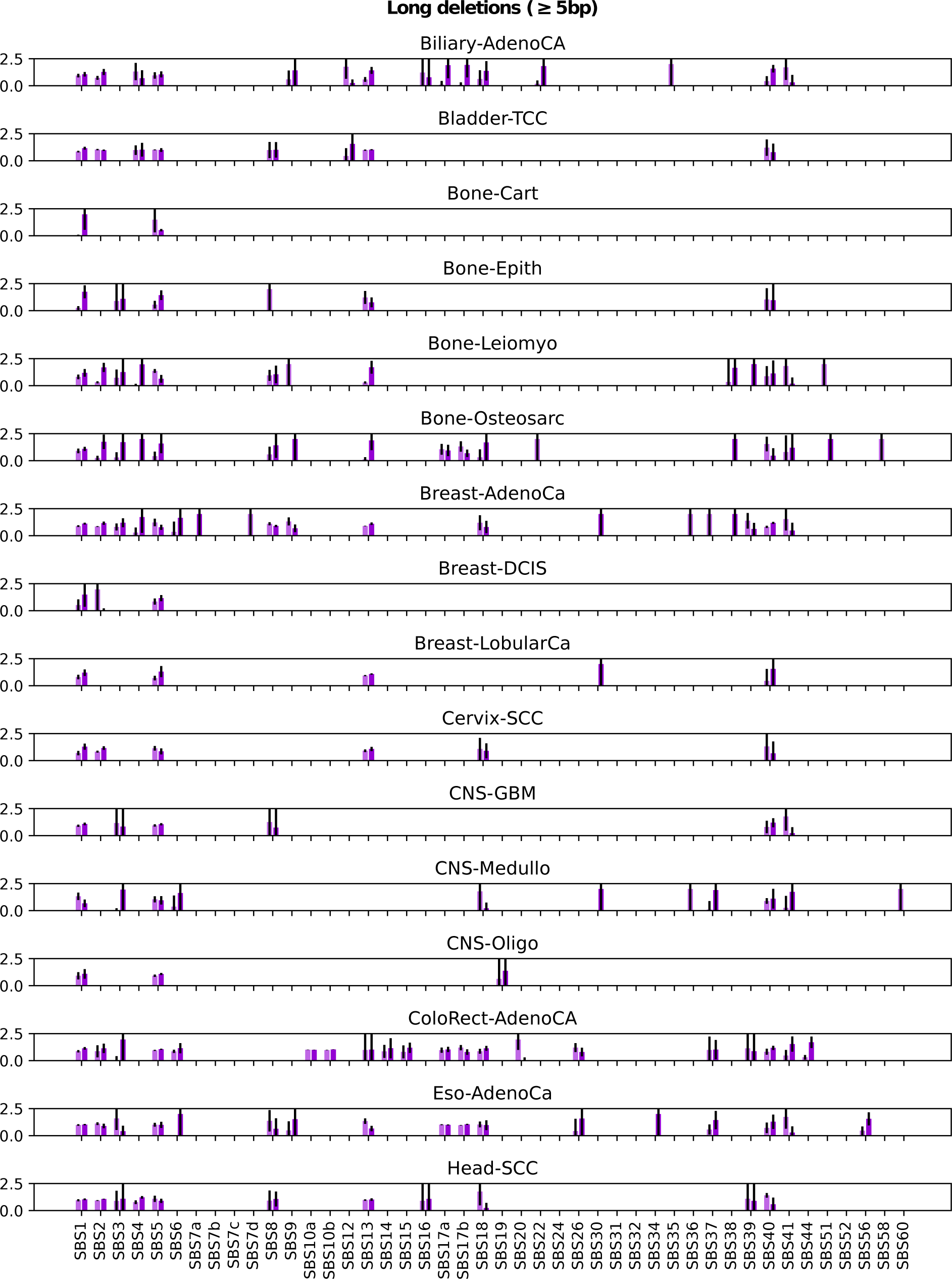
Scaled mutability of windows without long deletions (lighter color) and windows with long deletions (darker color) for all signatures for the first half of all cancer types in PCAWG. Error bars are computed through 100 bootstrap replicates.

**Extended Data Figure 14:**
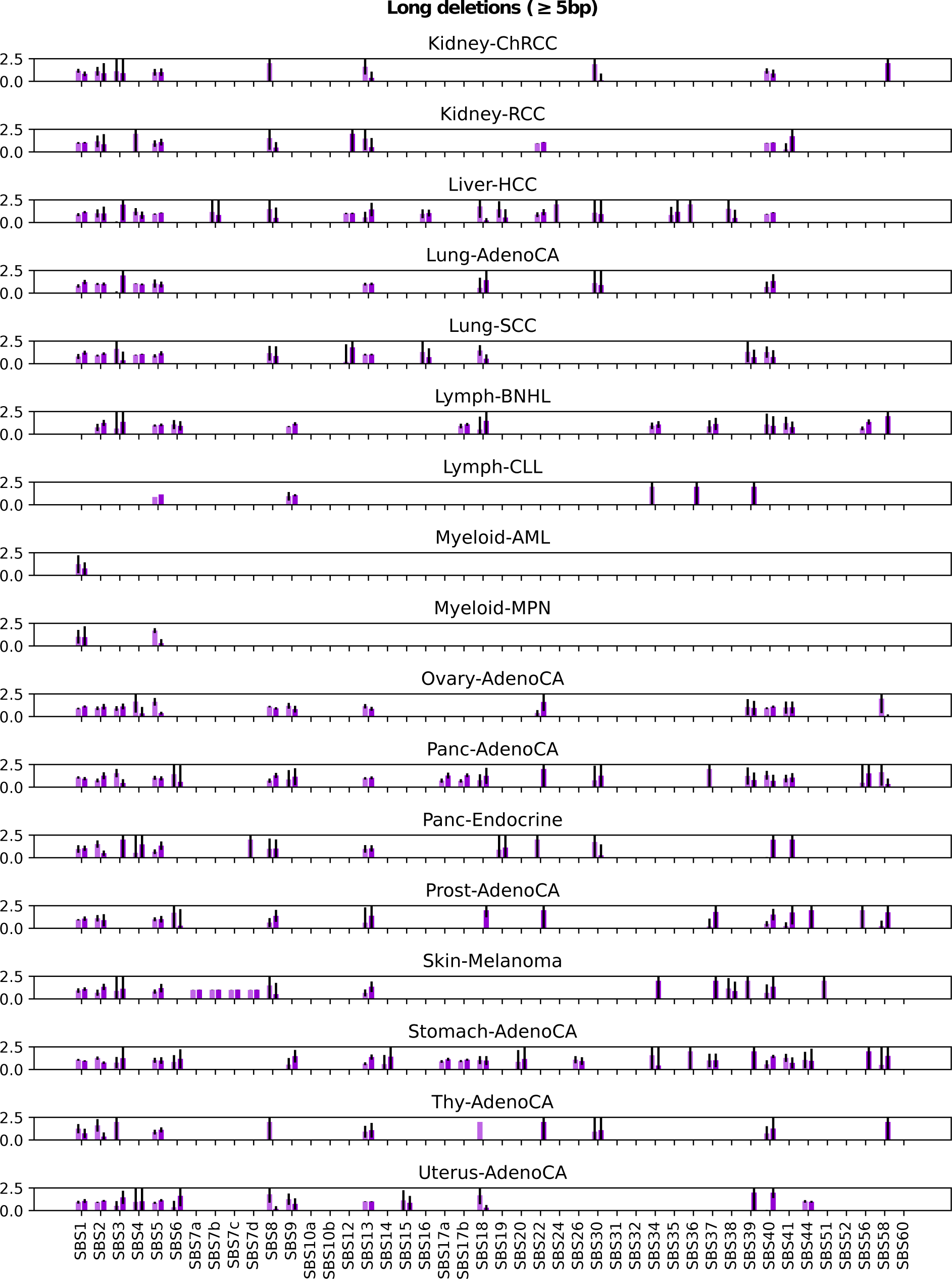
Scaled mutability of windows without long deletions (lighter color) and windows with long deletions (darker color) for all signatures for the second half of all cancer types in PCAWG. Error bars are computed through 100 bootstrap replicates.

**Extended Data Figure 15:**
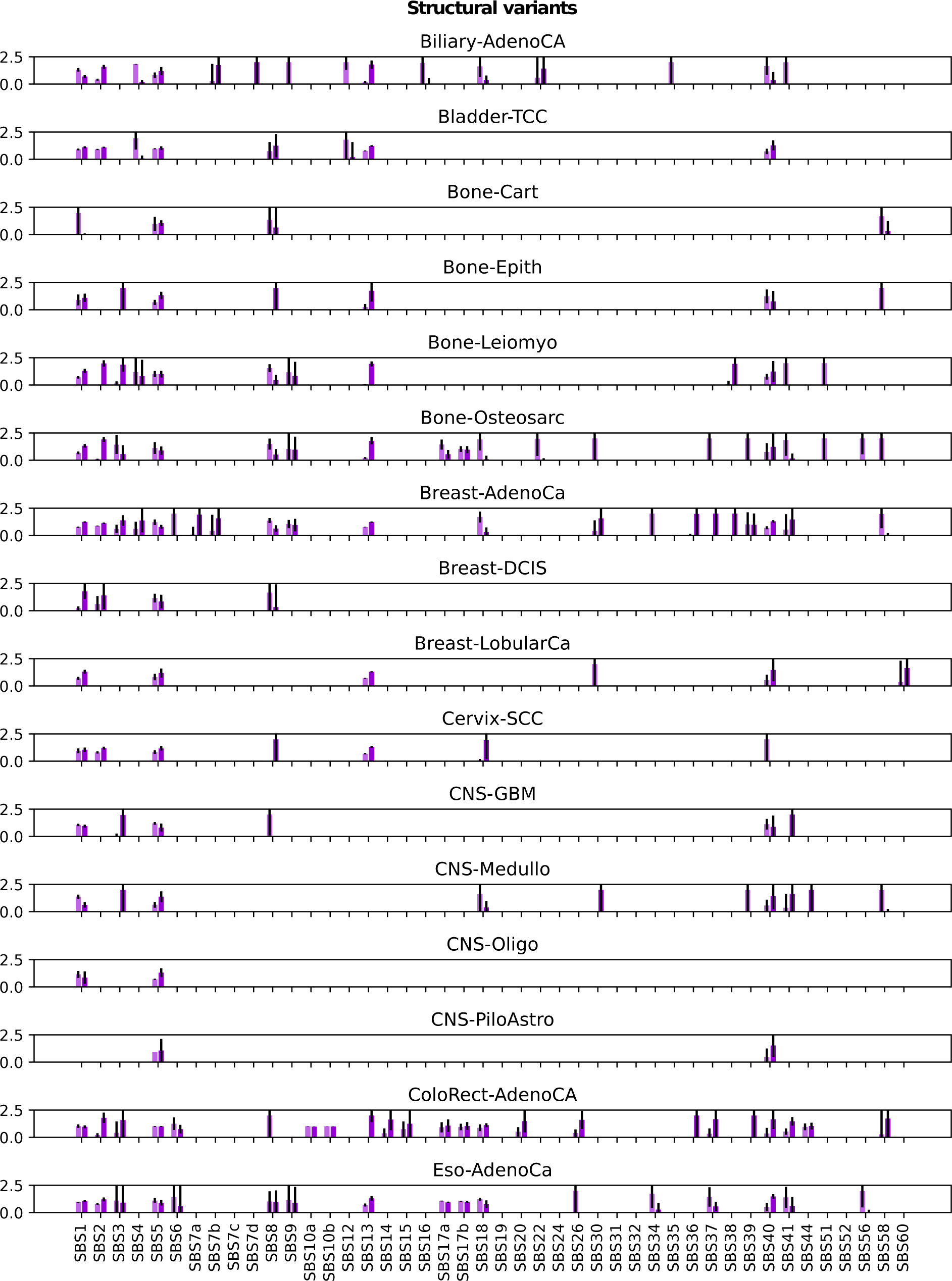
Scaled mutability of windows without structural variants (lighter color) and windows with structural variants (darker color) for all signatures for the first half of all cancer types in PCAWG. Error bars are computed through 100 bootstrap replicates.

**Extended Data Figure 16:**
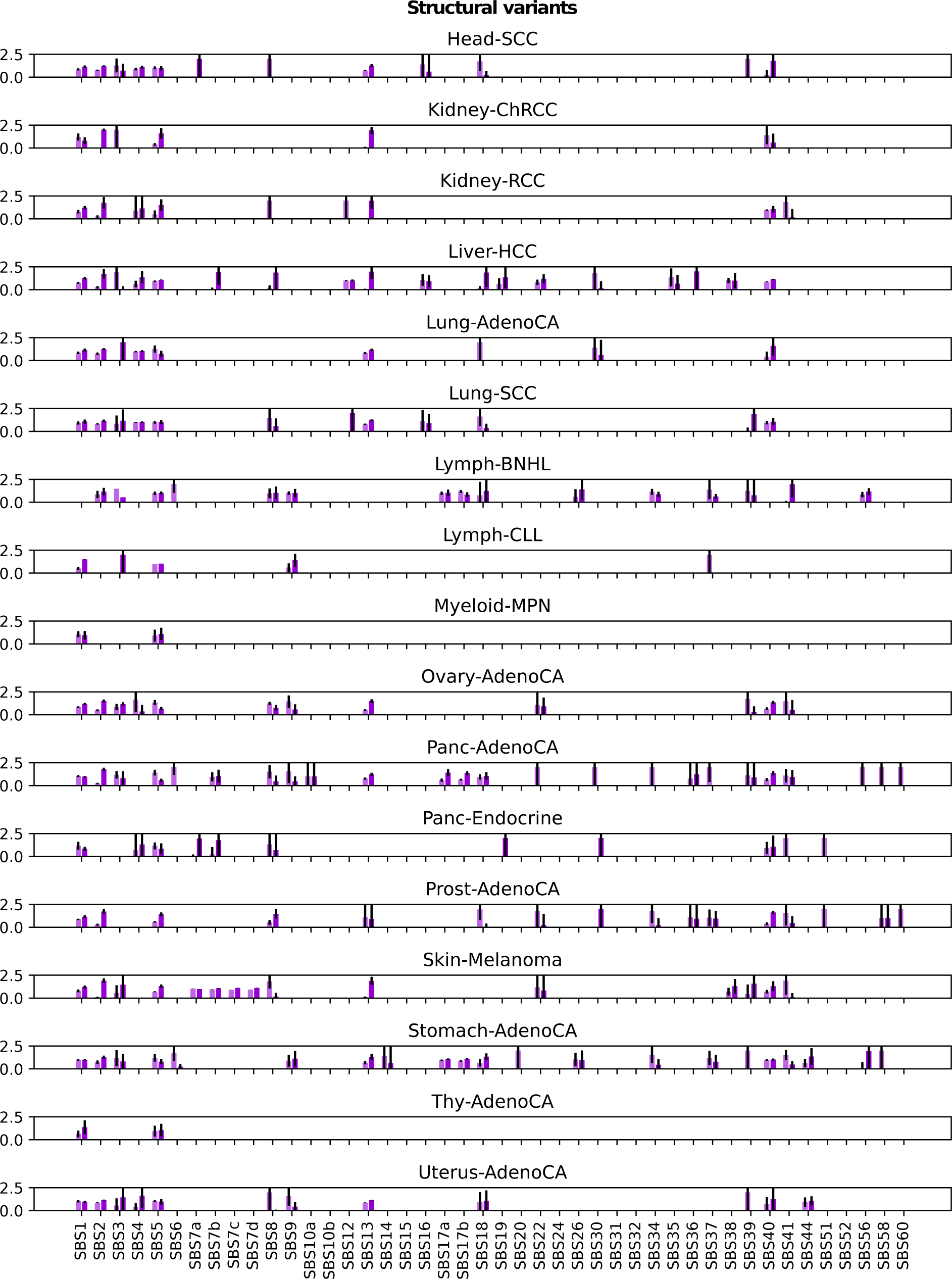
Scaled mutability of windows without structural variants (lighter color) and windows with structural variants (darker color) for all signatures for the second half of all cancer types in PCAWG. Error bars are computed through 100 bootstrap replicates.

**Extended Data Figure 17:**
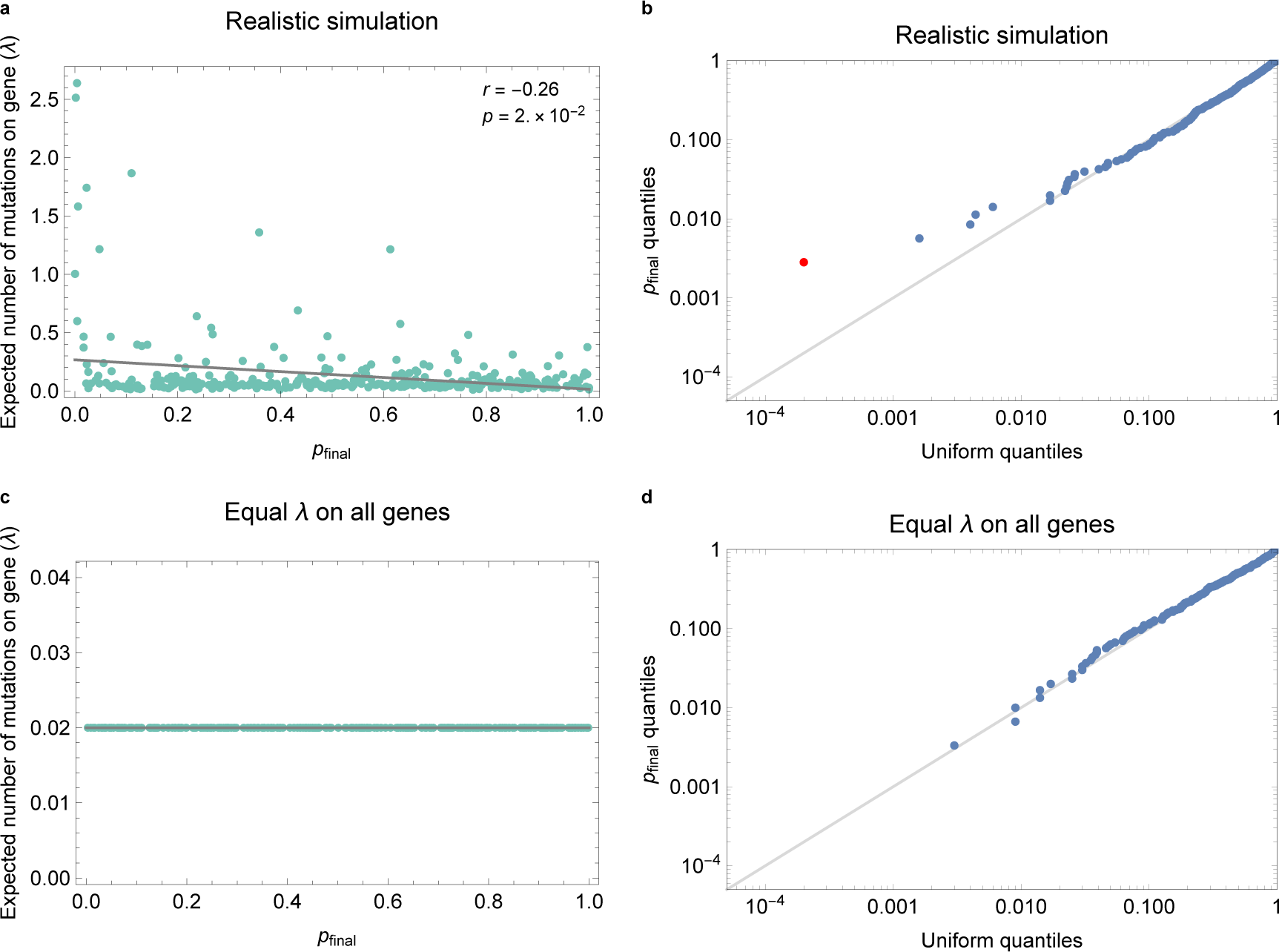
Results for simulations of 18000 genes and 130 tumours (mimicking the bladder cancer cohort in Kim *et al.* [69]). Genes have Poisson-distributed mutation counts with varying expected values, given by the parameter *λ*, which is drawn from an inverse gamma distribution with mean 0.005, compatible with common distributions inferred in cancer cohorts [105]. Ten genes were modelled as highly mutable genes by increasing their *λ* parameter by a factor *∈* [10, 130], approximately corresponding to the length ratio between the coding sequence of *TTN* and the average gene or a realistic driver gene selection strength (*≈* dN/dS). Tumour-specific average mutation probabilities were modulated with a log-normal-distributed random variable with mean 1.3. Like in Kim *et al.*, only genes mutated in at least 5% of patients were used, giving the same order of magnitude of used genes (10^2^). (a) Correlation between the expected number of mutations on the gene, *λ*, and its final p-value. The final p-value is obtained from the fraction of shuffled binary matrices, as output by the curveball algorithm [90], for which the gene had a p-value smaller than or equal to its real p-value based on a Mann-Whitney test of total mutation burden in tumours with a mutation in the gene *vs* those without. (b) Quantile-quantile plot of the distribution of final p-values against a uniform distribution. Statistically significant genes (after Benjamini-Hochberg correction) with FDR *≤* 0.1 are colored red. (c) By construction, the correlation between *λ* and the final p-value is zero when simulating a constant Poisson parameters for all genes. (d) Quantile-quantile plot for a simulation with constant Poisson parameter *λ* per gene and no tumour-specific modulation of the mutation probability.

## Notes

### Competing Interest Statement

The authors have declared no competing interest.

## References

[1] Schuster-Böckler, B. & Lehner, B. Chromatin organization is a major influence on regional mutation rates in human cancer cells. Nature 488, 504–507 (2012).

[2] Lawrence, M. S. et al. Mutational heterogeneity in cancer and the search for new cancer-associated genes. Nature 499, 214–218 (2013).

[3] Supek, F. & Lehner, B. Scales and mechanisms of somatic mutation rate variation across the human genome. DNA repair 81, 102647 (2019).

[4] Seplyarskiy, V. B. et al. Population sequencing data reveal a compendium of mutational processes in the human germ line. Science 373, 1030–1035 (2021).

[5] Cosmic mutational signatures (v3.3 - june 2022). https://cancer.sanger.ac.uk/signatures/sbs/. Accessed: 2022-08-24.

[6] Alexandrov, L. B. et al. The repertoire of mutational signatures in human cancer. Nature 578, 94–101 (2020).

[7] Rosenthal, R., McGranahan, N., Herrero, J., Taylor, B. S. & Swanton, C. deconstructSigs: Delineating mutational processes in single tumors distinguishes DNA repair deficiencies and patterns of carcinoma evolution. Genome Biology 17, 31 (2016).

[8] Hübschmann, D., et al. Analysis of mutational signatures with yet another package for signature analysis. Genes Chromosomes and Cancer 60, 314–331 (2021).

[9] Nik-Zainal, S. et al. Mutational Processes Molding the Genomes of 21 Breast Cancers. Cell 149, 979–993 (2012).

[10] Vali-Pour, M., Lehner, B. & Supek, F. The impact of rare germline variants on human somatic mutation processes. Nature communications 13, 1–21 (2022).

[11] Andrianova, M. A., et al. Extended family with germline pathogenic variant in polymerase delta provides strong evidence for recessive effect of proofreading inactivation. bioRxiv (2022).

[12] Pleasance, E. D. et al. A comprehensive catalogue of somatic mutations from a human cancer genome. Nature 463, 191–196 (2010).

[13] Alexandrov, L. B. et al. Mutational signatures associated with tobacco smoking in human cancer. Science (New York, N.Y.) 354, 618–622 (2016).

[14] Gurjao, C. et al. Discovery and features of an alkylating signature in colorectal cancer. Cancer discovery 11, 2446 (2021).

[15] Nik-Zainal, S. et al. Landscape of somatic mutations in 560 breast cancer whole genome sequences. Nature 534, 47–54 (2016).

[16] Farmer, H. et al. Targeting the DNA repair defect in BRCA mutant cells as a therapeutic strategy. Nature 434, 917–921 (2005).

[17] Bryant, H. E. et al. Specific killing of BRCA2-deficient tumours with inhibitors of poly (ADP-ribose) polymerase. Nature 434, 913–917 (2005).

[18] Brady, S. W., Gout, A. M. & Zhang, J. Therapeutic and prognostic insights from the analysis of cancer mutational signatures. Trends in Genetics (2021).

[19] Levatíc, J., Salvadores, M., Fuster-Tormo, F. & Supek, F. Mutational signatures are markers of drug sensitivity of cancer cells. Nature communications 13, 1–19 (2022).

[20] Morganella, S. et al. The topography of mutational processes in breast cancer genomes. Nature Communications 7, 11383 (2016).

[21] Vöhringer, H., Van Hoeck, A., Cuppen, E. & Gerstung, M. Learning mutational signatures and their multidimensional genomic properties with tensorsignatures. Nature Communications 12, 1–16 (2021).

[22] Hanahan, D. & Weinberg, R. A. Hallmarks of cancer: the next generation. Cell 144, 646–674 (2011).

[23] Kaplan, A. R. & Glazer, P. M. Impact of hypoxia on DNA repair and genome integrity. Mutagenesis 35, 61–68 (2020).

[24] Bhandari, V., Li, C. H., Bristow, R. G. & Boutros, P. C. Divergent mutational processes distinguish hypoxic and normoxic tumours. Nature Communications 11, 1–10 (2020).

[25] Ellrott, K. et al. Scalable open science approach for mutation calling of tumor exomes using multiple genomic pipelines. Cell systems 6, 271–281 (2018).

[26] The ICGC/TCGA Pan-Cancer Analysis of Whole Genomes Consortium. Pan-cancer analysis of whole genomes. Nature 578, 82–93 (2020).

[27] Rozen, S. G. & Yang, W. Synthetic (mutational) signature generation (synsiggen). https://github.com/steverozen/SynSigGen (2022).

[28] Krüger, S. & Piro, R. M. DecompTumor2Sig: Identification of mutational signatures active in individual tumors. BMC Bioinformatics 20, 1–15 (2019).

[29] Blokzijl, F., Janssen, R., van Boxtel, R. & Cuppen, E. MutationalPatterns: Comprehensive genome-wide analysis of mutational processes. Genome Medicine 10, 1–11 (2018).

[30] Fantini, D., Vidimar, V., Yu, Y., Condello, S. & Meeks, J. J. MutSignatures: an R package for extraction and analysis of cancer mutational signatures. Scientific Reports 2020 10:1 10, 1–12 (2020).

[31] Huang, X., Wojtowicz, D. & Przytycka, T. M. Detecting presence of mutational signatures in cancer with confidence. Bioinformatics 34, 330 (2018).

[32] Omichessan, H., Severi, G. & Perduca, V. Computational tools to detect signatures of mutational processes in DNA from tumours: A review and empirical comparison of performance. PLoS ONE 14, e0221235 (2019).

[33] Lynch, A. Decomposition of mutational context signatures using quadratic programming methods. F1000Research 5, 1253 (2016).

[34] Li, S., Crawford, F. W. & Gerstein, M. B. Using sigLASSO to optimize cancer mutation signatures jointly with sampling likelihood. Nature communications 11, 3575 (2020).

[35] Jin, H. et al. Accurate and sensitive mutational signature analysis with musical. bioRxiv 2022–04 (2022).

[36] Priestley, P. et al. Pan-cancer whole-genome analyses of metastatic solid tumours. Nature 575, 210–216 (2019).

[37] Moore, L. et al. The mutational landscape of human somatic and germline cells. Nature 597, 381–386 (2021).

[38] Yizhak, K. et al. RNA sequence analysis reveals macroscopic somatic clonal expansion across normal tissues. Science 364, eaaw0726 (2019).

[39] Blokzijl, F. et al. Tissue-specific mutation accumulation in human adult stem cells during life. Nature 538, 260–264 (2016).

[40] Blanc, V. & Davidson, N. O. C-to-U RNA Editing: Mechanisms Leading to Genetic Diversity. Journal of Biological Chemistry 278, 1395–1398 (2003).

[41] Kiran, A. & Baranov, P. V. DARNED: a DAtabase of RNa EDiting in humans. Bioinformatics 26, 1772–1776 (2010).

[42] Hanahan, D. & Weinberg, R. A. The hallmarks of cancer. Cell 100, 57–70 (2000).

[43] Luoto, K. R., Kumareswaran, R. & Bristow, R. G. Tumor hypoxia as a driving force in genetic instability. Genome Integrity 4, 5 (2013).

[44] Grin, I. & Ishchenko, A. A. An interplay of the base excision repair and mismatch repair pathways in active DNA demethylation. Nucleic acids research 44, 3713–3727 (2016).

[45] Poulos, R. C., Olivier, J. & Wong, J. W. The interaction between cytosine methylation and processes of dna replication and repair shape the mutational landscape of cancer genomes. Nucleic acids research 45, 7786–7795 (2017).

[46] Meier, B. et al. Mutational signatures of DNA mismatch repair deficiency in C. elegans and human cancers. Genome research 28, 666–675 (2018).

[47] Seplyarskiy, V. B. & Sunyaev, S. The origin of human mutation in light of genomic data. Nature Reviews Genetics 1–15 (2021).

[48] Sanders, M. A. et al. Life without mismatch repair. bioRxiv (2021).

[49] Gelova, S. P., Doherty, K. N., Alasmar, S. & Chan, K. Intrinsic base substitution patterns in diverse species reveal links to cancer and metabolism. Genetics 222, iyac144 (2022).

[50] Gelova, S. P. & Chan, K. Mutagenesis induced by protonation of single-stranded dna is linked to glycolytic sugar metabolism. Mutation Research/Fundamental and Molecular Mechanisms of Mutagenesis 826, 111814 (2023).

[51] Hammond, E. M., Dorie, M. J. & Giaccia, A. J. Inhibition of ATR leads to increased sensitivity to hypoxia/reoxygenation. Cancer research 64, 6556–6562 (2004).

[52] Denko, N. C. Hypoxia, HIF1 and glucose metabolism in the solid tumour. Nature Reviews Cancer 8, 705–713 (2008).

[53] Daijo, H. et al. Cigarette smoke reversibly activates hypoxia-inducible factor 1 in a reactive oxygen species-dependent manner. Scientific Reports 6, 34424 (2016).

[54] Buffa, F. M., Harris, A. L., West, C. M. & Miller, C. J. Large meta-analysis of multiple cancers reveals a common, compact and highly prognostic hypoxia metagene. British Journal of Cancer 102, 428–435 (2010).

[55] Ragnum, H. B. et al. The tumour hypoxia marker pimonidazole reflects a transcriptional programme associated with aggressive prostate cancer. British Journal of Cancer 112, 382– 390 (2015).

[56] Winter, S. C. et al. Relation of a hypoxia metagene derived from head and neck cancer to prognosis of multiple cancers. Cancer Research 67, 3441–3449 (2007).

[57] Velasco-Hernandez, T., et al. A comprehensive single-cell expression atlas of human AML leukemia-initiating cells unravels the contribution of HIF pathway and its therapeutic potential (2022).

[58] Davies, H. et al. HRDetect is a predictor of BRCA1 and BRCA2 deficiency based on mutational signatures. Nature medicine 23, 517–525 (2017).

[59] Otlu, B., et al. Topography of mutational signatures in human cancer. Cell Reports 42 (2023).

[60] Vöhringer, H., Hoeck, A. V., Cuppen, E. & Gerstung, M. Learning mutational signatures and their multidimensional genomic properties with tensorsignatures. Nature Communications 2021 12:1 12, 1–16 (2021).

[61] Gerstung, M. et al. The evolutionary history of 2,658 cancers. Nature 578, 122–128 (2020).

[62] Rubanova, Y. et al. Reconstructing evolutionary trajectories of mutation signature activities in cancer using TrackSig. Nature communications 11, 1–12 (2020).

[63] Abécassis, J., Reyal, F. & Vert, J.-P. Clonesig can jointly infer intra-tumor heterogeneity and mutational signature activity in bulk tumor sequencing data. Nature communications 12, 1–16 (2021).

[64] Shinde, J. et al. Palimpsest: an R package for studying mutational and structural variant signatures along clonal evolution in cancer. Bioinformatics 34, 3380–3381 (2018).

[65] McGranahan, N. et al. Clonal status of actionable driver events and the timing of mutational processes in cancer evolution. Science translational medicine 7, 283ra54–283ra54 (2015).

[66] Dentro, S. C. et al. Characterizing genetic intra-tumor heterogeneity across 2,658 human cancer genomes. Cell 184, 2239–2254 (2021).

[67] Christensen, S. et al. 5-fluorouracil treatment induces characteristic t¿ g mutations in human cancer. Nature communications 10, 4571 (2019).

[68] Perillo, B. et al. ROS in cancer therapy: The bright side of the moon. Experimental & molecular medicine 52, 192–203 (2020).

[69] Kim, J. et al. Somatic ercc2 mutations are associated with a distinct genomic signature in urothelial tumors. Nature genetics 48, 600–606 (2016).

[70] Spisak, N., de Manuel, M., Milligan, W. R., Sella, G. & Przeworski, M. Disentangling sources of clock-like mutations in germline and soma. bioRxiv 2023–09 (2023).

[71] Fricker, M., et al. Chronic cigarette smoke exposure induces systemic hypoxia that drives intestinal dysfunction. JCI insight 3 (2018).

[72] Thompson, P. S. & Cortez, D. New insights into abasic site repair and tolerance. DNA repair 90, 102866 (2020).

[73] Nik-Zainal, S. et al. Landscape of somatic mutations in 560 breast cancer whole-genome sequences. Nature 534, 47–54 (2016).

[74] Kolinjivadi, A. M. et al. Moonlighting at replication forks–a new life for homologous recombination proteins brca 1, brca 2 and rad 51. FEBS letters 591, 1083–1100 (2017).

[75] Mateos-Gomez, P. A. et al. Mammalian polymerase *θ* promotes alternative nhej and suppresses recombination. Nature 518, 254–257 (2015).

[76] Ceccaldi, R. et al. Homologous-recombination-deficient tumours are dependent on pol*θ*-mediated repair. Nature 518, 258–262 (2015).

[77] Wyatt, D. W. et al. Essential roles for polymerase *θ*-mediated end joining in the repair of chromosome breaks. Molecular cell 63, 662–673 (2016).

[78] Schrempf, A., et al. Pol*θ* processes ssdna gaps and promotes replication fork progression in brca1-deficient cells. Cell Reports 41 (2022).

[79] Belan, O. et al. Polq seals post-replicative ssdna gaps to maintain genome stability in brca-deficient cancer cells. Molecular cell 82, 4664–4680 (2022).

[80] Seki, M. et al. High-efficiency bypass of dna damage by human dna polymerase q. The EMBO journal 23, 4484–4494 (2004).

[81] Caron, P., Pobega, E. & Polo, S. E. Dna double-strand break repair: all roads lead to heterochromatin marks. Frontiers in Genetics 12, 730696 (2021).

[82] Gong, F. & Miller, K. M. Histone methylation and the dna damage response. Mutation Research/Reviews in Mutation Research 780, 37–47 (2019).

[83] Gursoy-Yuzugullu, O., Ayrapetov, M. K. & Price, B. D. Histone chaperone anp32e removes h2a. z from dna double-strand breaks and promotes nucleosome reorganization and dna repair. Proceedings of the National Academy of Sciences 112, 7507–7512 (2015).

[84] Alatwi, H. E. & Downs, J. A. Removal of h2a. z by ino 80 promotes homologous recombination. EMBO reports 16, 986–994 (2015).

[85] van Schendel, R., van Heteren, J., Welten, R. & Tijsterman, M. Genomic scars generated by polymerase theta reveal the versatile mechanism of alternative end-joining. PLoS genetics 12, e1006368 (2016).

[86] Feng, W. et al. Genetic determinants of cellular addiction to dna polymerase theta. Nature communications 10, 4286 (2019).

[87] Brambati, A., Barry, R. M. & Sfeir, A. Dna polymerase theta (pol*θ*)–an error-prone polymerase necessary for genome stability. Current opinion in genetics & development 60, 119–126 (2020).

[88] Petljak, M. et al. Mechanisms of apobec3 mutagenesis in human cancer cells. Nature 607, 799–807 (2022).

[89] Hogg, M., Seki, M., Wood, R. D., Doublíe, S. & Wallace, S. S. Lesion bypass activity of dna polymerase *θ* (polq) is an intrinsic property of the pol domain and depends on unique sequence inserts. Journal of molecular biology 405, 642–652 (2011).

[90] Strona, G., Nappo, D., Boccacci, F., Fattorini, S. & San-Miguel-Ayanz, J. A fast and unbiased procedure to randomize ecological binary matrices with fixed row and column totals. Nature communications 5, 4114 (2014).

[91] Taglialatela, A. et al. Rev1-pol*ζ* maintains the viability of homologous recombination-deficient cancer cells through mutagenic repair of primpol-dependent ssdna gaps. Molecular Cell 81, 4008–4025 (2021).

[92] McVey, M., Khodaverdian, V. Y., Meyer, D., Cerqueira, P. G. & Heyer, W.-D. Eukaryotic dna polymerases in homologous recombination. Annual review of genetics 50, 393–421 (2016).

[93] Kotsantis, P., Petermann, E. & Boulton, S. J. Mechanisms of oncogene-induced replication stress: jigsaw falling into place. Cancer discovery 8, 537–555 (2018).

[94] Gonźalez-Olvera, J. C., et al. Protonation of nucleobases in single-and double-stranded dna. ChemBioChem 19, 2088–2098 (2018).

[95] Grosse, N. et al. Deficiency in homologous recombination renders mammalian cells more sensitive to proton versus photon irradiation. International Journal of Radiation Oncology* Biology* Physics 88, 175–181 (2014).

[96] Wood, R. D. & Doublíe, S. Dna polymerase *θ* (polq), double-strand break repair, and cancer. DNA repair 44, 22–32 (2016).

[97] Goodfellow, I., Bengio, Y. & Courville, A. Deep Learning (MIT Press, 2016). http://www.deeplearningbook.org.

[98] Maas, A. L., Hannun, A. Y. & Ng, A. Y. Rectifier nonlinearities improve neural network acoustic models. In ICML Workshop on Deep Learning for Audio, Speech and Language Processing (2013).

[99] Kingma, D. P. & Ba, J. Adam: A method for stochastic optimization. In Bengio, Y. & LeCun, Y. (eds.) 3rd International Conference on Learning Representations, ICLR 2015, San Diego, CA, USA, May 7-9, 2015, Conference Track Proceedings (2015).

[100] Chen, D. & Plemmons, R. J. Nonnegativity constraints in numerical analysis. In A. Bultheel and R. Cools (Eds.), Symposium on the Birth of Numerical Analysis, World Scientific (Press, 2009).

[101] Pearce, T., Brintrup, A., Zaki, M. & Neely, A. High-quality prediction intervals for deep learning: A distribution-free, ensembled approach. In International conference on machine learning, 4075–4084 (PMLR, 2018).

[102] Kingma, D. P. & Welling, M. Auto-encoding variational Bayes (2014). 1312.6114.

[103] Kingma, D. P., Salimans, T. & Welling, M. Variational dropout and the local reparameterization trick (2015).

[104] Smith, T. C. A., Arndt, P. F. & Eyre-Walker, A. Large scale variation in the rate of germ-line de novo mutation, base composition, divergence and diversity in humans. PLoS Genetics 14, e1007254 (2018). URL https://www.ncbi.nlm.nih.gov/pmc/articles/PMC5891062/.

[105] Weghorn, D. & Sunyaev, S. Bayesian inference of negative and positive selection in human cancers. Nature Genetics 49, 1785 (2017).

