## Supplementary Tables 1-15 for "Mutational signature decomposition with deep neural networks reveals origins of clock-like processes and hypoxia dependencies": README.txt

Fig 2:
supplementary_table_1.csv: MC3 SigNet decompositions and classification.
supplementary_table_2.csv: HMF SigNet decompositions and classification.
supplementary_table_3.csv: Moore SigNet decompositions and classification.
supplementary_table_4.csv: GTEx SigNet decompositions and classification.

Fig 3:
supplementary_table_5.csv: list of genes for each hypoxia score used. 
supplementary_table_6.csv: data for hypoxia analysis using PCAWG dataset. 
supplementary_table_7.csv: data for hypoxia analysis using MC3 dataset.

Fig 4:
supplementary_table_8.csv: data for WG/WE.
supplementary_table_9.csv: data for Clonal/Subclonal.

Fig 5:
supplementary_table_10.csv: regression coefficients replication time.
supplementary_table_11.csv: histones p-values
supplementary_table_12.csv: replication time decompositions.
supplementary_table_13.csv: short indels decompositions.
supplementary_table_14.csv: long deletions decompositions.
supplementary_table_15.csv: structural variants decompositions.

 
